# Gut Microbiota predicts Healthy Late-life Aging in Male Mice

**DOI:** 10.1101/2021.06.22.449472

**Authors:** Shanlin Ke, Sarah J. Mitchell, Michael R. MacArthur, Alice E. Kane, David A. Sinclair, Emily M. Venable, Katia S. Chadaideh, Rachel N. Carmody, Francine Grodstein, James R. Mitchell, Yang-Yu Liu

**Affiliations:** Channing Division of Network Medicine, Brigham and Women’s Hospital and Harvard Medical School, Boston, Massachusetts 02115, USA; State Key Laboratory of Pig Genetic Improvement and Production Technology, Jiangxi Agricultural University 330045, China; Department of Molecular Metabolism, Harvard T.H. Chan School of Public Health, Boston, MA, 02115, USA; Department of Health Sciences and Technology, ETH Zurich, Zurich 8005 Switzerland; Blavatnik Institute, Dept. of Genetics, Paul F. Glenn Center for Biology of Aging Research at Harvard Medical School, Boston, MA 02115 USA; Department of Human Evolutionary Biology, Harvard University, Cambridge, MA, 02138, USA; Department of Epidemiology, Harvard T.H. Chan School of Public Health, Boston, MA, 02115, USA

**Author notes:** To whom correspondence should be addressed: Y.-Y.L. and S.J.M.

## Abstract

Calorie restriction (CR) extends lifespan and retards age-related chronic diseases in most species. There is growing evidence that the gut microbiota has a pivotal role in host health and age-related pathological conditions. Yet, it is still unclear how CR and the gut microbiota are related to healthy aging. Here we report findings from a small longitudinal study of male C57BL/6 mice maintained on either *ad libitum* or mild (15%) CR diets from 21 months of age and tracked until natural death. We demonstrate that CR results in a significant reduction in frailty index (FI), a well-established indicator of aging. We observed significant alterations in bacterial load, diversity, and compositional patterns of the mouse gut microbiota during the aging process. Interrogating the FI-related microbial features using machine learning techniques, we show that gut microbial signatures from 21-month-old mice can predict the healthy aging of 30-month-old mice with reasonable accuracy. This study deepens our understanding of the links between CR, gut microbiota, and frailty in the aging process of mice.

## Introduction

The proportional population of older persons is growing across the globe^1^. This demographic shift will increase the prevalence of age-related disease and place a significant burden on health costs and social care. Moreover, increased longevity (i.e., lifespan) does not necessarily translate to better quality of life (i.e., healthspan)^2^. Thus, it is imperative to improve our understanding of mechanisms underlying aging processes and develop practical interventions to promote healthy aging and delay age-related diseases.

Aging is one of the most complex biological processes that affects a wide array of physiological, genomic, metabolic, and immunological functions^9,10^. These age-related functional changes can lead to organ and systemic decline, which ultimately results in death. There is now growing evidence that the gut microbiota interacts with these physiological functions, and thereby plays a pivotal role in host health and age-related pathological conditions^3–5^. The gut microbiota is regulated by a complex interplay between host and environmental factors, including age, diet, antibiotics, genetics, and lifestyle^7,8^. In turn, changes in the gut microbiota can alter host physiology, increasing the incidence and/or severity of many diseases that contribute to morbidity and mortality in later life, such as inflammatory bowel disease^17^, type 2 diabetes^19^, obesity^20^, cardiovascular disease^21^, and neurodegenerative disease^22^. During host aging, the gut microbiota undergoes dramatic changes in composition and function^12–16^. The gut microbiota of elderly people is different from that of adults^14,23,24^, and microbial compositions in the elderly correlate with measures of frailty, barrier dysfunction, gut motility, and inflammation^25^. Nevertheless, the extent to which these changes result from host aging or contribute to it remains unclear. Unlike other organs, the gut microbiota might not be expected to follow the same general trajectory of somatic senescence^11^.

Calorie restriction (CR), a dietary regimen that reduces the consumption of food without resulting in malnutrition, has been shown in animal models to retard development of age-related chronic diseases and extend the lifespan^26–29^. In addition to effects on host physiology, CR can also reshape the gut microbial community in both humans^30,31^ and animal models^32–34^. CR-induced alterations to the gut microbiome might play a role in extending lifespan and healthspan and delaying the onset of age-related disorders. In this study, we evaluate how the gut microbiota changes during the aging process in mice and test whether gut microbial features can predict healthy aging. To do this, we performed quantitative PCR (qPCR) targeting the 16S rRNA gene and 16S rRNA gene sequencing of bacterial DNA extracted from fecal samples from a cohort of aging male mice tracked from 21 months of age. We investigated associations between these microbial signatures and biomarkers of host condition, including weight, food intake, hematological markers, and frailty index (FI), a validated biomarker of biological age that is a strong predictor of mortality, morbidity, and other age-related outcomes^35^. Examining how signatures in the gut microbiota predict future aging status can illuminate the utility of the gut microbiota as an early indicator of healthy aging.

## Results

### Experimental design

The experimental design is shown in Fig. 1. Following baseline phenotypic measurements (body weight, food intake, frailty index, grip strength, and fecal collection), adult male C57BL/6 mice were randomized at 21 months of age into *ad libitum* diet (AL, n=14) or mild calorie restriction diet (CR, 15% fewer calories than their peers consuming an *ad libitum* diet, n=8) groups and followed longitudinally until death. From each birth cohort that we received, we randomized the mice equally into groups to avoid a strong birth-cohort effect. We repeated phenotypic measurements after 9 months (30 months of age) and recorded survival. We performed qPCR targeting the 16S rRNA gene as well as 16S rRNA gene sequencing on 44 stool samples, collected at 21 and 30 months of age, from 22 mice.

**Fig. 1.**
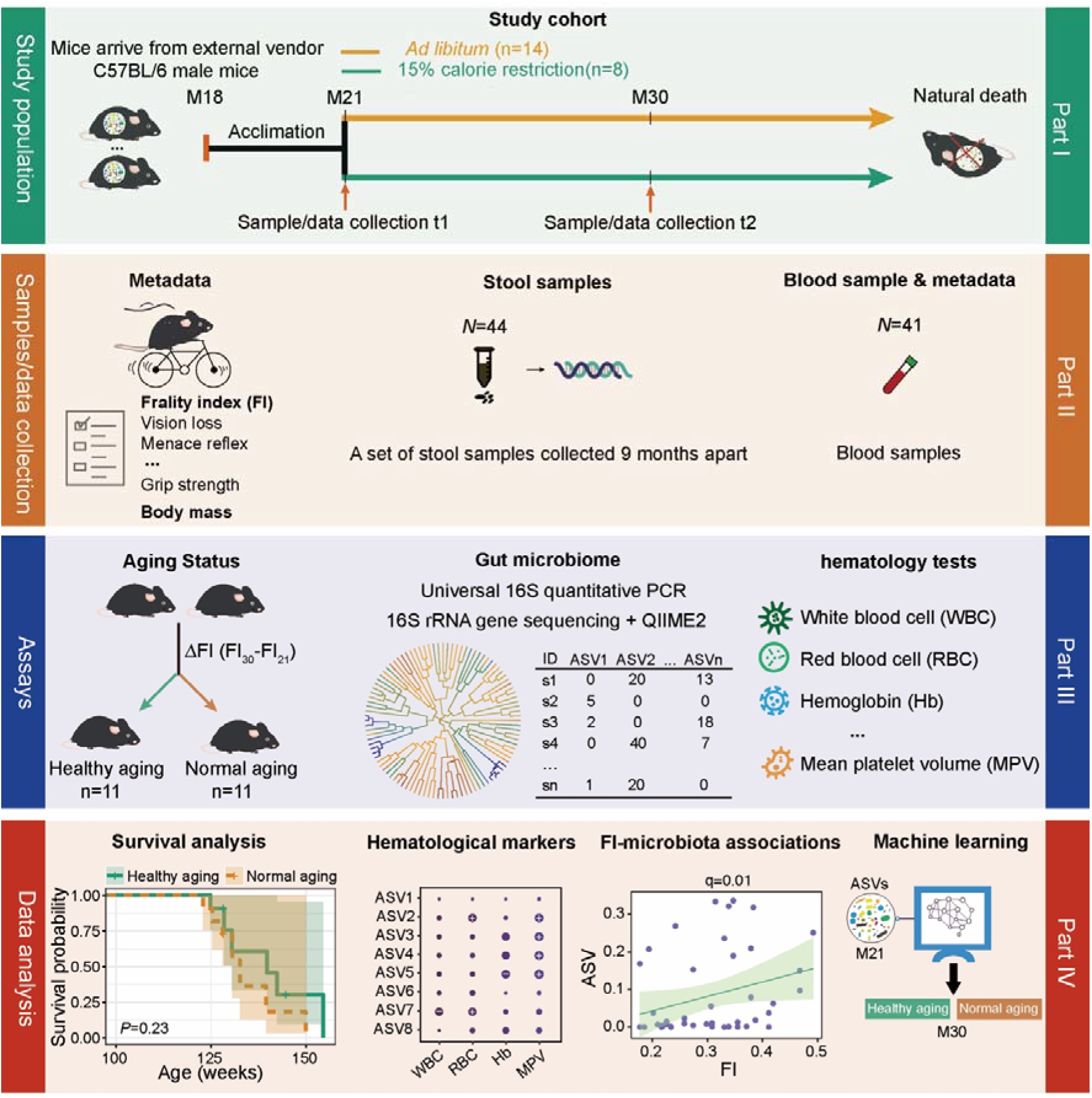
Schematic diagram showing the experimental design. The study cohort was comprised of 22 adult male C57BL/6 mice, which were recruited into the study at 21 months of age after having been maintained since birth under standard husbandry conditions (see Methods). We collected blood and fecal samples and measured frailty using a compound index at 21 months (baseline) and 30 months of age. Following baseline measurements, we randomly divided these mice into two diet groups, fed either *ad libitum* (AL, n=14) with standard chow or under mild (15%) calorie restriction (CR, n=8). Mice were then followed longitudinally until death. We performed universal 16S quantitative PCR (qPCR) to quantify absolute bacterial abundance and used QIIME2 to obtain the ASV microbial features. Blood markers were measured using standard methods. We then used the median FI change (denoted as ΔFI) between 21 and 30 months of age to delineate healthy versus normal aging.

### The association of the physiological characteristics with chronological age

The mouse clinical frailty index (FI) is based on established clinical signs of deterioration in mice^36,37^. Briefly, the clinical assessment includes evaluation of the integument, the musculoskeletal system, the vestibulocochlear/auditory systems, ocular and nasal systems, digestive system, urogenital system, respiratory system, signs of discomfort, body weight, and body surface temperature. FI score is continuous from 0-1, with higher values indicating worse frailty. A cutoff of 0.21 has been previously used in rodents^38^ to stratify frailty as either high (frail: FI≥0.21) or low (not frail: FI<0.21). But as mice reaches 30 months old, they all become frail with higher FI score (FI>0.21) in our study. Indeed, as shown in Fig. 2a and Fig. S1a, FI score significantly increased with chronological age from 21 to 30 months at the population level (*P*-value = 4.8e-06, Wilcoxon signed-rank test). Hence, instead of using a fixed FI score cutoff, in this work we used the median value of FI change (denoted as ΔFI) to delineate healthy versus normal aging. Specifically, we calculated ΔFI between month 21 and 30 for each mouse, and then we dichotomized those mice at month 30 into two groups based on the medium value of their ΔFI: ‘healthy aging’ (age in weeks: mean 121.78□± standard deviation 3.88; ΔFI: 0.088□±□0.038; FI: 0.342□±□0.048; n=11); and ‘normal aging’ (age in weeks: 121.42□±□4.07; ΔFI: 0.179□±□0.034; FI: 0.398□±□0.055; n=11). CR diet was associated with a lower level of ΔFI at month 30 than AL diet (Fig. 2b, *P*-value = 0.029, Wilcoxon–Mann–Whitney test). In particular, 87.5% (7/8) of mice with CR diet belonged to the healthy aging group compared to just 36.4% (4/11) of mice fed *ad libitum*. These results suggest that CR had a beneficial effect on aging, consistent with previous studies^27^.

**Fig. 2.**
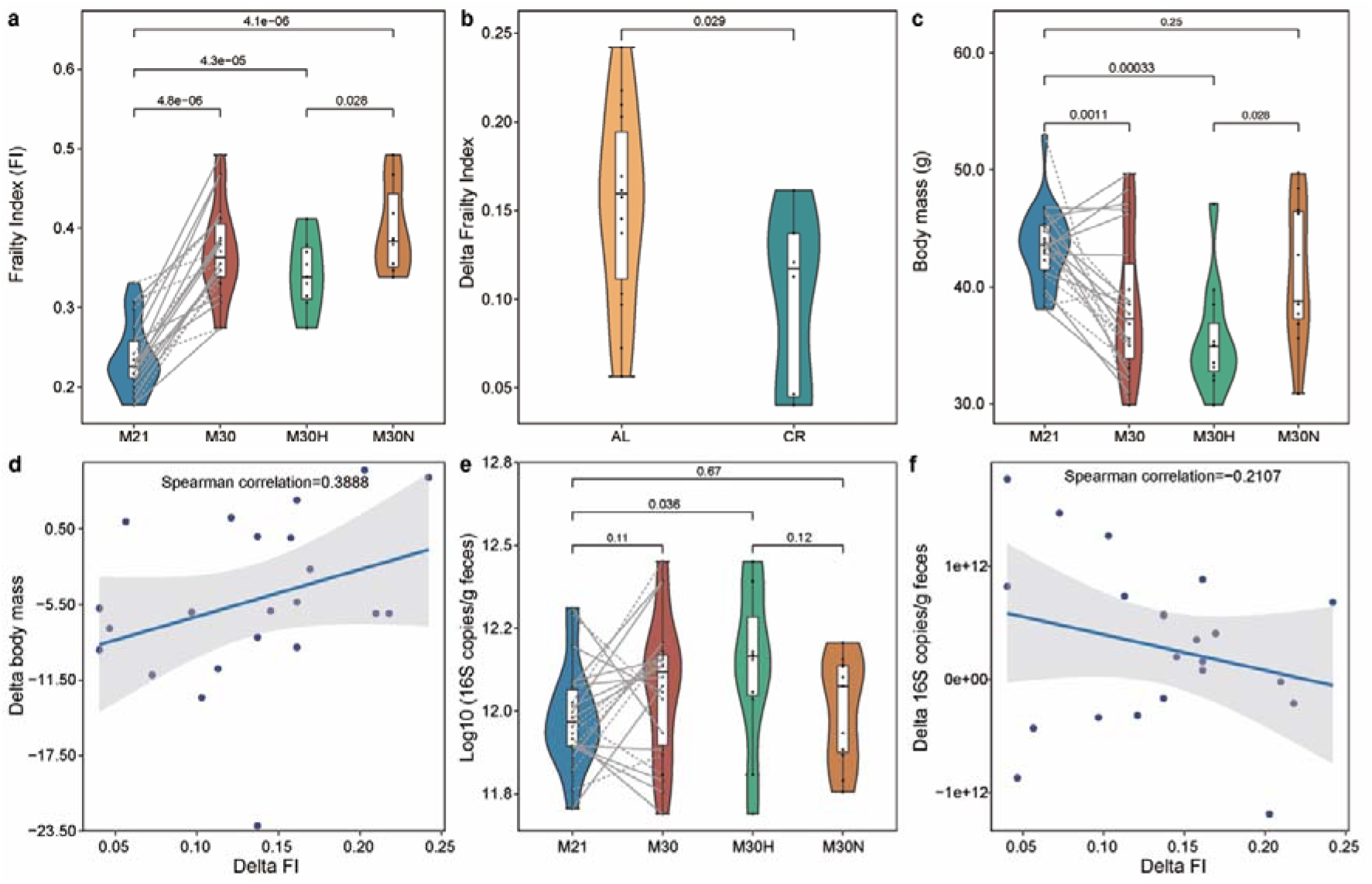
Frailty index associates with chronological age in mice. **a**, Frailty index changes with age. Mice at 30 months of age were grouped into healthy and normal aging based on the median ΔFI. **b**, The effect of caloric restriction on the ΔFI between 21 and 30 months of age. **c**, Comparison of body mass (BM) for different groups. **d**, The association between ΔFI and ΔBM in all mice. **e**, Comparison of total bacterial load for different groups. **f**, The association between ΔFI and ΔBL in all mice. Points obtained for the same subject from 21 and 30 months of age are joined by solid (AL diet) and dotted (CR diet) lines. *P* value shown in **a**-**c** and **e** are the result of Wilcoxon–Mann–Whitney test (unpaired) and Wilcoxon signed rank test (paired). The correlation coefficient shown in **d** and **f** is the result of Spearman correlation. The lines show lm fit for the data, and shaded areas show 95% confidence intervals for the fit.

We found that the body mass (BM) of mice generally decreased during aging (Fig. 2c, *P*-value = 0.0011, Wilcoxon signed-rank test), which was contributed by healthy aging mice due to the fact that most of them (63.64%) were from the CR group (Fig. S1b). At 30 months of age, the BM of the healthy aging mice was significantly lower than the normal aging (Fig. 2c, *P*-value = 0.028, Wilcoxon–Mann–Whitney test) and baseline mice (Fig. S1b, *P*-value = 0.0049, Wilcoxon signed-rank test). To better understand this finding, we calculated delta change of BM (ΔBM) between month 21 and 30 for each mouse. The ΔFI was positively associated with ΔBM (Fig. 2d, *ρ* = 0.3888, Spearman correlation), suggesting that a normal aging mouse (with large ΔFI) is associated with an increasing level of BM. In addition, we found that the BM in healthy aging mice gradually decreased over time (Fig. S2a), especially in those mice with CR diet (Fig. S2b). Additionally, normal aging mice showed rapid loss of BM after some time points (Fig. S2). Using Kaplan–Meier survival analysis, the differences in cumulative survival rates were not statistically significant between healthy and normal aging mice (Fig. S3, *P*-value = 0.23, log-rank test). However, the healthy aging mice showed qualitatively longer lifespan (134.36 ± 9.43) than normal aging (131.06 ± 7.53) mice (*P*-value = 0.313, Wilcoxon–Mann–Whitney test), as some mice from the healthy aging group lived substantially longer.

### Aging-related changes in gut microbial community

Using universal 16S qPCR, we first measured the total bacterial load (BL) in the stool samples (Fig. 2e and Fig S1c). The results showed the total BL detected in these healthy aging mice was higher than the BL present in the normal aging mice (Fig. 2e). For the changes of total BL over time (ΔBL), we found ΔFI was inversely associated with ΔBL (Fig. 2f, *ρ* = −0.2107, Spearman correlation), suggesting that a normal aging mouse (larger ΔFI) is associated with a decreasing total BL.

We then measured the gut microbial community compositions of those stool samples using 16S rRNA gene sequencing (see Methods, Table S1). Phylum-level taxonomic profiles of the gut microbiome samples of those mice are shown in Fig. 3a. Consistent with previous studies^39,40^, we found that Bacteroidetes, Firmicutes and Verrucomicrobia were the most dominant phyla in the murine gut microbiota. Notable age-related compositional shifts included an enrichment in Firmicutes, and reduction in Bacteroidetes and Verrucomicrobia, although such trade-offs among dominant phyla are expected *a priori* in relative abundance data. Moreover, the Firmicutes/Bacteroidetes ratio of the gut microbiota increased with age (Fig. 3b, *P*-value = 0.0025, Wilcoxon signed-rank test). Both healthy aging and normal aging mice showed higher values for this ratio compared with baseline mice (Fig. S4a).

**Fig. 3.**
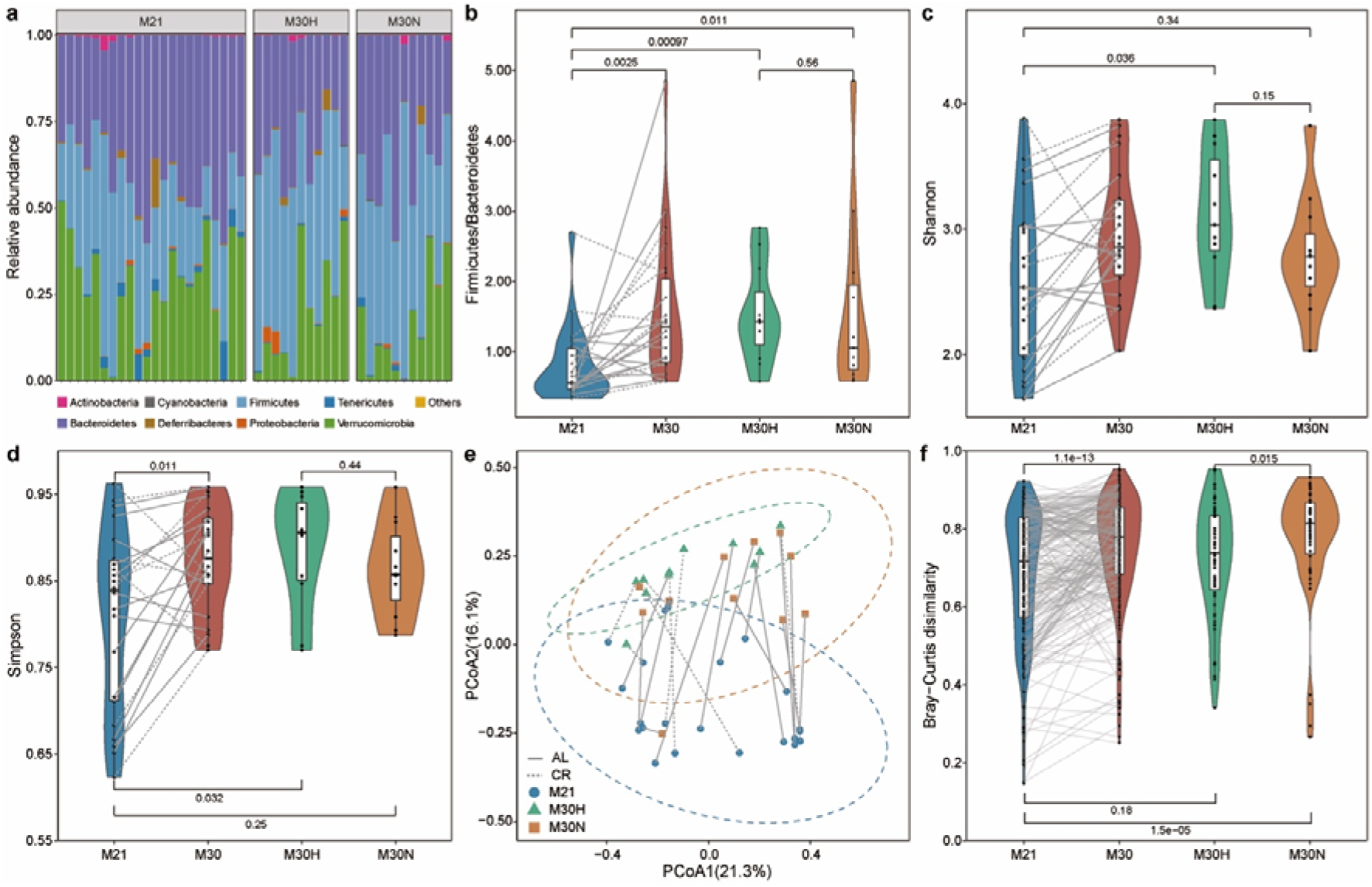
Impact of aging on gut microbial communities. **a**, Relative abundance of bacterial phyla. **b**, The ratio of Firmicutes to Bacteroidetes. Alpha diversity using Shannon (**c**) and Simpson (**d**) index. **e**, Beta diversity using Principal Coordinate Analysis (PCoA) of Bray–Curtis dissimilarity. The dotted ellipse borders with color represent the 95% confidence interval. **f**, Boxplot of gut microbiota Bray–Curtis dissimilarity between subjects within each group. Points obtained for the same subject from 21 and 30 months of age in **b**-**e** are joined by solid (AL diet) and dotted (CR diet) lines. Points obtained for the same subject pairs from 21 and 30 months of age in **f** are joined by solid line. *P* value shown in **b**-**d**, and **f** are the result of Wilcoxon–Mann–Whitney test (unpaired) and Wilcoxon signed rank test (paired).

Using the Shannon diversity and Simpson index as alpha diversity measures, we found that alpha diversity increased with age (Fig. 3c,d and Fig. S4b,c), consistent with a previous mouse study^41^. Interestingly, we found that the Shannon diversity was only significantly higher in healthy aging mice compared to baseline mice (Fig. S4b, *P*-value = 0.019, Wilcoxon signed-rank test). In addition, a clear separation (permutational multivariate analysis of variance (PERMANOVA) test, *P*-value = 0.0001, Bray-Curtis dissimilarity) could be seen between mice at 21 and 30 months of age in the principal coordinate analysis (PCoA) plot based on Bray-Curtis dissimilarity (Fig. 3e). Indeed, PERMANOVA test indicated significantly altered microbial compositions for both healthy aging (*P*-value = 0.0004) and normal (*P*-value = 0.0086) aging mice between baseline and 30 months of age (Fig. S4d). However, we found no significant difference between healthy aging and normal aging mice at both 21 (*P*-value = 0.8747) and 30 (*P*-value = 0.3536) months of age. Bray-Curtis dissimilarity was higher among individuals within normal aging mice compared to baseline mice (Fig. S4e, *P*-value = 4e-08, Wilcoxon signed-rank test) or healthy aging mice (Fig. 3f, *P*-value = 0.015, Wilcoxon–Mann–Whitney test). This suggests that normal aging is characterized by high variations in gut microbiota between individuals.

### The effect of aging on hematology and associations between gut microbiota and blood markers

Aging is associated with a decline in immune system function at multiple levels^42^. To explore aging-related immune system modifications, we measured hematological parameters over time (Table S2). We found that the mice at 30 months of age tended to have higher level (with *P* value <0.05) of neutrophils percentage, neutrophil to lymphocyte ratio (NLR), monocytes percentage (MOp, % of leukocytes), red cell distribution width (RDW, % variation), and mean platelet volume (MPV, fL), but lower level (with *P* value <0.05) of white blood cell (WBC, k/uL), lymphocytes (LY, k/uL), lymphocytes percentage (LYp, % of leukocytes), red blood cell (RBC, M/uL), hemoglobin (Hb, g/dL), Mean corpuscular volume (MCV, fL) and hematocrit (HCT, % volume) when compared with mice at 21 months of age. Specifically, higher NLR (an important biomarker of systemic inflammation^43^) levels on 30-month-old mice were mainly observed in normal aging mice (*P* value = 0.016). Here *P* values were all calculated from the Wilcoxon–Mann–Whitney test, adjusted with the Benjamini–Hochberg FDR method. These results confirm prior observations that high levels of inflammation are not an inevitable consequence of aging, but rather associated with normal or unhealthy aging. Moreover, at 30 months of age, we found that normal aging mice had significantly higher MPV but normal PLT.

Given the effects of aging process on hematology, we next used MaAsLin2 (multivariate analysis by linear models^44^) to evaluate the associations between microbial taxa and blood markers. These linear mixed models accounted for within-individual correlation from the study’s repeated sampling design, as well as occasional missing observations at some time points. To control for potential confounding variables, we added four covariates into the model as fixed effects, including diet treatment, cohort, cage, and body mass. In addition, each mouse’s identifier treated as random effect. A total of 24 ASVs (amplicon sequence variant) features were significantly associated with at least one blood marker (Fig. 4, *q*-value ≤ 0.2, Table S3). In general, blood markers correlating most with microbial taxa included MCV, LY and NLR. For example, MCV was inversely associated with the abundance of ASV 3949 (*Anaerotruncus, q*-value = 2.38e-14) and ASV3729 (*Clostridium aldenense, q*-value = 1.52e-6), and LY was positively associated with ASV890 (Ruminococcaceae, *q*-value = 0.0004), ASV2868 (*Oscillibacter, q*-value = 0.015), and ASV2973 (*Intestinimonas butyriciproducens, q*-value = 0.035). NLR was positively associated with ASV5690 (*Flavonifractor plautii, q*-value = 0.04) and ASV555 (*Acetatifactor muris, q*-value = 0.048), and negatively associated with ASV2878 (Lachnospiraceae, *q*-value = 0.028), ASV4558 (Bacteroidales, *q*-value = 0.146), and ASV1970 (*Clostridium XlVa, q*-value = 0.189).

**Fig. 4.**
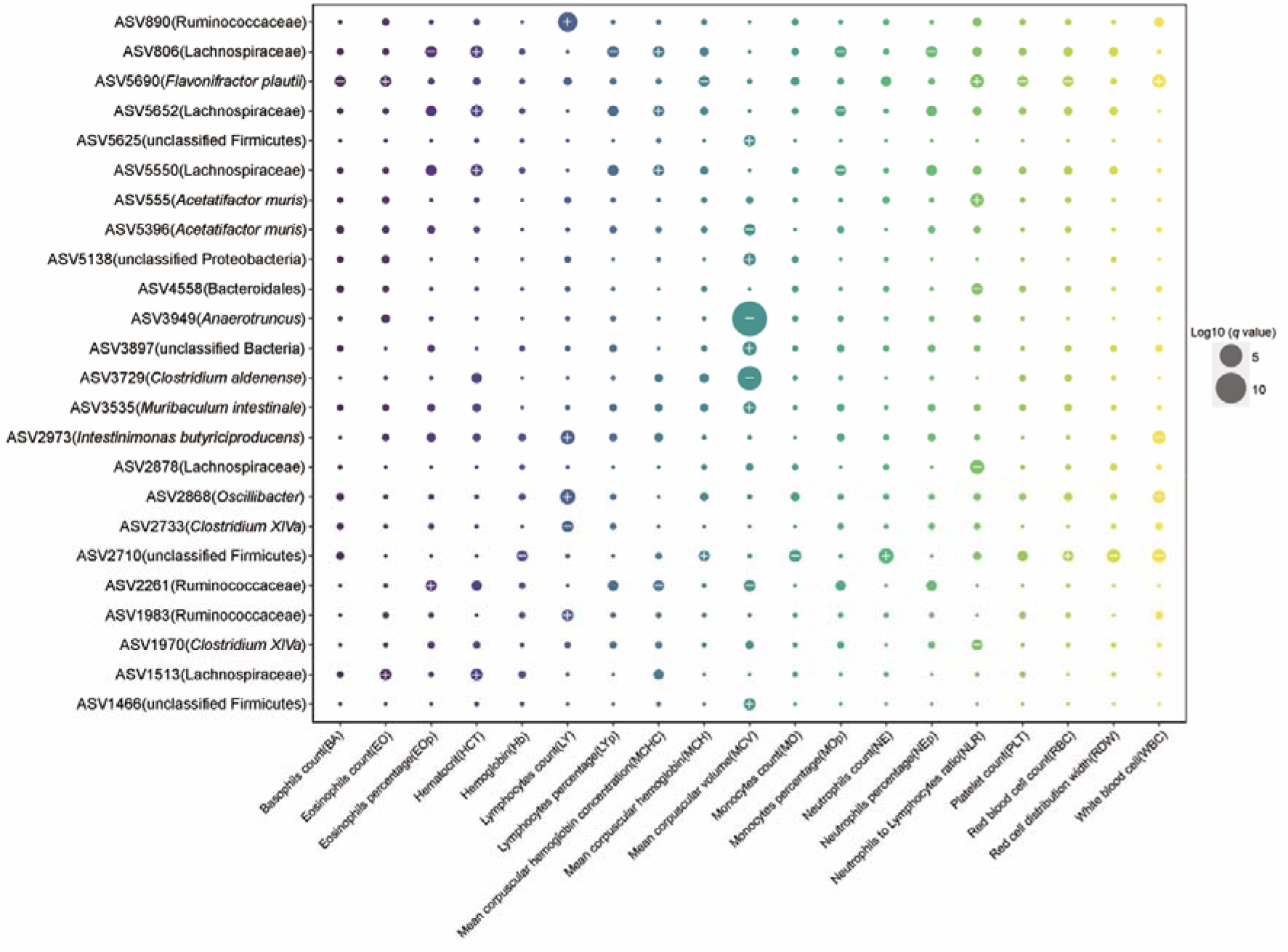
Identification of associations between blood cell and gut microbial features. Dot plot showing the links between the blood markers and gut microbial taxa identified using MaAsLin2. The sizes of dots represent the *q* values from MaAsLin2. The greater the size, the more significant the association. Symbols indicate the directions of associations in a given model: plus, significant positive associations; minus, significant negative associations. Threshold for FDR corrected *q*-value was set at 0.2. Linear mixed effects models were applied to the association with subject set as random-effect.

### Microbial taxa related to frailty index and healthy aging

We next investigated the FI in relation to the microbial features using MaAsLin2 in which diet, cohort, cage, and body mass were included as fixed effects and each mouse’s identifier was included as a random effect. We observed a set of 14 microbial features that were strongly linked to FI (Fig. 5, *q-*value ≤ 0.2, Table S4). Consistent with previous reports that the abundance of the *Clostridium sensu stricto* genus increases with aging^45–47^, ASV3100 (*Clostridium sensu stricto: q*-value = 0.021) was positively associated with the FI. *Clostridium XlVa*^48^ (ASV2882, *q*-value = 0.048 and ASV1101: *q*-value = 0.112) and *Subdoligranulum variabile*^49^ (ASV157, *q*-value = 0.153), known as important producers of butyrate, were found to be negatively associated with FI. We also found inverse associations of the FI with taxa such as ASV847 (*Phocea massiliensis, q*-value = 0.069), ASV 1726 (*Parabacteroides goldsteinii, q*-value = 0.083), and ASV1123 (*Enterorhabdus, q*-value = 0.090). A previous study linked *Parabacteroides goldsteinii* with reduction of intestinal inflammation and enhancement of cellular mitochondrial and ribosomal activities in the colon^50^.

**Fig. 5.**
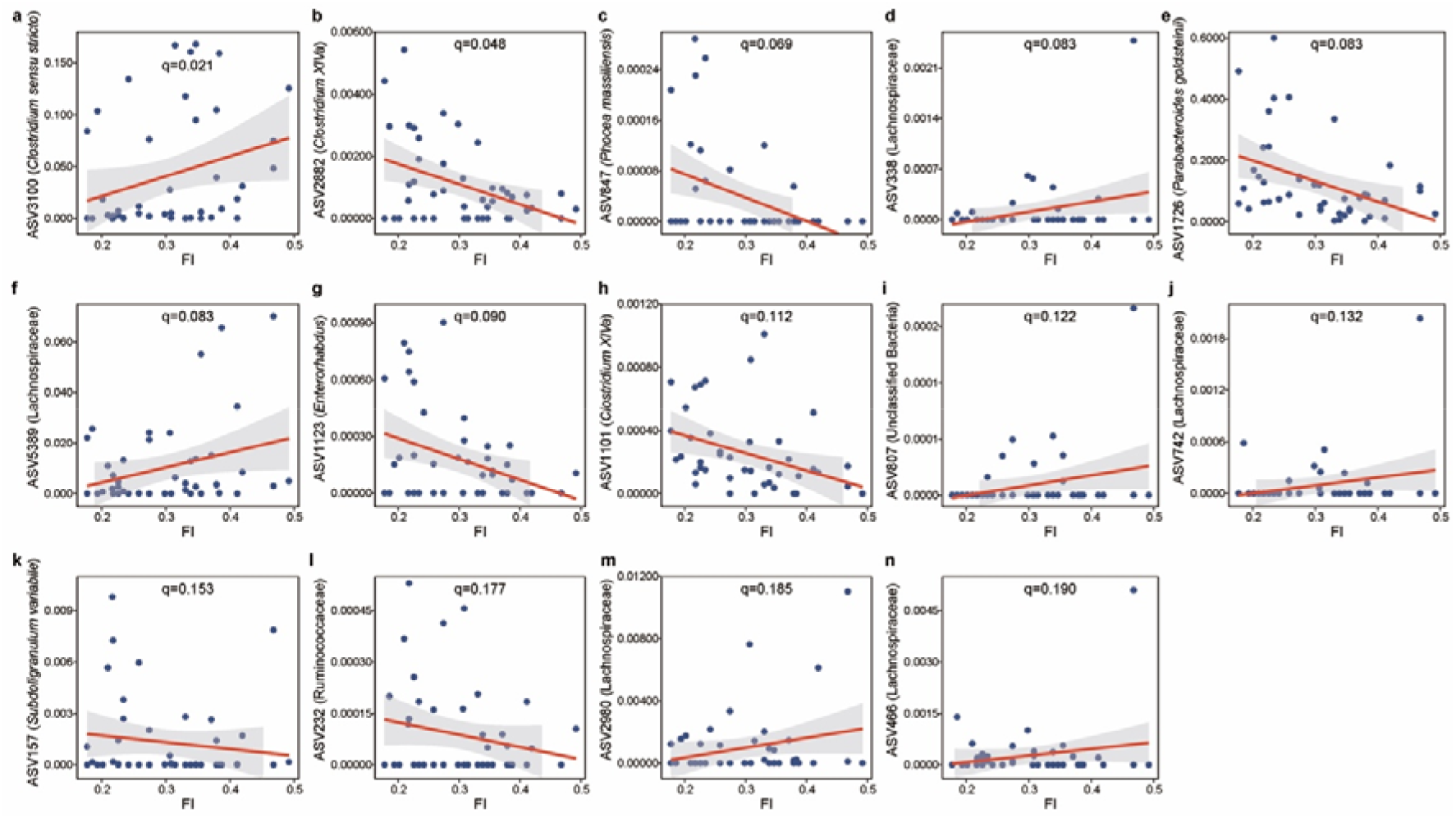
The significant associations between FI and gut microbial features. **a**, ASV3100 (*Clostridium sensu stricto*). **b**, ASV2882 (*Clostridium XlVa*). **c**, ASV847 (*Phocea massiliensis*). **d**, ASV338 (Lachnospiraceae). **e**, ASV1726 (*Parabacteroides goldsteinii*). **f**, ASV5389 (Lachnospiraceae). **g**, ASV1123 (*Enterorhabdus*). **h**, ASV1101 (*Clostridium XlVa*). **i**, ASV807 (Unclassified Bacteria). **j**, ASV742 (Lachnospiraceae). **k**, ASV157 (*Subdoligranulum variabile*). **l**, ASV232 (Ruminococcaceae). **m**, ASV2980 (Lachnospiraceae). **n**, ASV466 (Lachnospiraceae). Data shown are the relative abundance versus FI for ASVs that were significantly associated with FI in MaAsLin2. Threshold for FDR corrected *q*-value was set at 0.2. Linear mixed-effects models (LMMs) were applied to the association with subject set as random effect. The lines show lm fit for the data, and shaded areas show 95% confidence intervals for the fit.

To examine potential gut microbial signatures of late-life aging, we performed differential abundance analysis using ANCOM^51^ (analysis of composition of microbiomes). ANCOM identified multiple gut microbiota signature that were significantly different between baseline and 30 months of age in healthy aging (Fig. S5a and Table S5) and normal aging (Fig. S5b and Table S6) mice. Most of these features were also identified when comparing all mice between 21 and 30 months of age as a group (Fig. S6 and Table S7). Intriguingly, we found 7 ASVs that significantly and concordantly increased with age in both healthy aging and normal aging groups (Fig. S5), including ASV5550 (Lachnospiraceae), ASV5652 (Lachnospiraceae), ASV806 (Lachnospiraceae), ASV5435 (*Muribaculum intestinale*), ASV3224 (*Clostridium cocleatum*), ASV5628 (*Muribaculum intestinale*), and ASV3370 (*Muribaculum intestinale*), hinting at a universal murine microbial signature of aging. To assess how the microbial features links with healthy aging, we calculated the differential abundance of features between healthy aging and normal aging groups at both 21 and 30 months of age (Fig. S7). Our data found 6 (Fig. S7a, Table S8) and 9 (Fig. S7b, Table S9) ASVs were significantly associated with aging status at baseline and 30 months of ages, respectively. In particular, a set of microbial features were significantly enriched in healthy aging mice at 30 months of age, for example ASV648 (*Akkermansia muciniphila*), ASV73 (Ruminococcaceae), and ASV2756 (*Acetatifactor muris*). *A. muciniphila* has been observed previously to prevent the age-related decline in thickness of the colonic mucus layer and attenuate inflammation in old age^52^. Here, this microbial feature was detected and shown to be positively associated with healthy aging. Normal aging mice showed increased ASV3370 (*Muribaculum intestinale*), ASV3100 (*Clostridium sensu stricto*), ASV3939 (*Turicibacter sanguinis*), and ASV1123 (*Enterorhabdus*) compared with healthy aging mice. Consistent with positive relationship between FI and ASV3100 (*Clostridium sensu stricto*), we found that this feature was significantly higher in the normal aging group.

### Gut microbiota-based machine learning model to predict healthy aging

As microbial compositions were associated with aging status, we sought to determine whether the microbial features observed in mid-life could predict healthy aging in later life. To achieve that, we employed an Elastic-net (ENET) logistic regression model to predict healthy aging. Specifically, the ENET model trained with ASVs (present in at least 10% samples) achieved an accuracy of 0.5 (11/22) with leave-one-out cross-validation (LOOCV) (Fig. 6a). In principle, we can apply feature selection techniques to choose a subset of features from the dataset. However, to improve the biological meaning of the model, we then only selected the microbial features that significantly associated with FI. This approach included a microbial signature comprised of 14 ASVs (Fig. 6b) from the gut microbiota of 21-month old mice that exhibited power in predicting the healthy aging status of 30-month old mice with a LOOCV accuracy of 0.773 (17/22) (Fig. 6a). Notably, we also observed that *Clostridium sensu stricto* and *Enterorhabdus* were significantly overrepresented in normal aging mice at 30 months of age. A previous study found that *Clostridium sensu stricto* was significantly enriched in early onset necrotizing enterocolitis subjects^53^. *Enterorhabdus*, a member of the family Coriobacteriaceae, has been isolated from a mouse model of spontaneous colitis^54^. These findings were consistent with higher level of NLR in normal aging mice, which was used as a marker of systemic inflammation. This may partially explain the ability of these features to predict healthy aging over the subsequent 9 months. Finally, we validated our model by generating a null model with randomly selected features (number of features=14, times=100), which yielded a mean LOOCV accuracy of 0.443 (Fig. 6a).

**Fig. 6.**
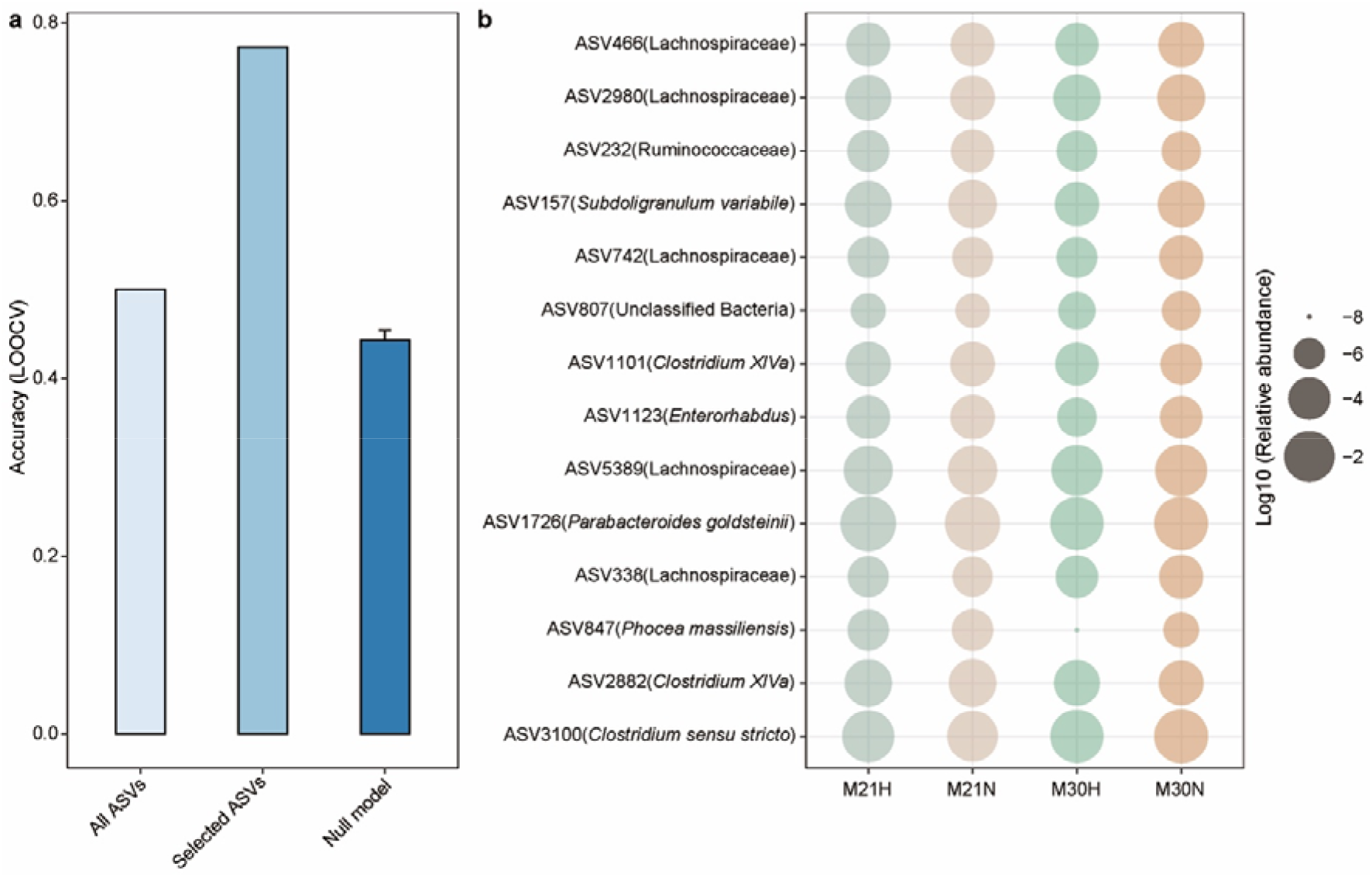
A gut microbiota-based signature moderately predicts healthy aging. **a,** Leave-one-out (LOOCV) accuracy evaluating ability to predict healthy aging using Elastic-net (ENET). Each bar represents the performance based on different microbial feature combination: all ASVs, 14 FI-associated ASVs, and null model with 14 randomly selected features run 100 times. **b,** The mean relative abundance of 14 FI-related ASVs across different groups. The healthy aging status at 21 months of age was determined by the aging status at 30 months of age. Relative abundances are plotted on log10 scale. Error bars represent the standard errors of the means (SEM) in null model.

## Discussion

Over the last few decades, global average life expectancy has increased dramatically, resulting in a proportionately larger aging population. Currently, chronological age is the most widely used indictor of aging, yet it provides limited information on the quality of life during the aging process. Understanding how to promote healthy aging will be key to increasing healthspan. Evidence is emerging that the gut microbiota is intrinsically linked with energy metabolism and the aging process^55–58^. In this study, we observed that the mouse gut microbiota is associated with healthy aging on late-life aged mice. And we identified a specific stool-microbiota-derived signature of aging that yielded a reasonable accuracy for the prediction of healthy aging.

A better predictor of mortality and morbidity in humans than chronological age is the Frailty index (FI)^61^. The FI has been reverse translated into a tool for mice which includes 31 non-invasive parameters across a range of systems^37,62^. Previous studies applied 0.21 as a cut-off point of FI to stratify between high frailty (≥0.21) or low frailty (<0.21)^38,63,64^. Given this specific threshold provides limited insight into the aging process, we instead employed the ΔFI (FI changes between 30 and 21 months of age) to quantify the ability to maintain health conditions during aging. Indeed, those mice with higher ΔFI (based on median value) were more vulnerable and frail. In our study, we only included the mice with basic measurements and biological samples at both 21 and 30 months, resulting 22 male mice that were fed either AL (n=14) or CR (n=8) diets. To avoid the issue arising from imbalanced sample size, we stratified the mice to healthy and normal aging mice based on the ΔFI. As expected, 87.5% (7/8) of mice with CR diet belonged to the healthy aging group compared to just 36.4% (4/11) of mice fed AL.

Although several previous studies demonstrated the links between gut microbiota and aging in mice, these studies mainly focused on the comparison between different growth stages^65–67^. In this study, we examined the gut microbiota collected at 21 and 30 months of age from 22 mice and measured the aging status. Concordant with previous reports, we found that aging was associated with increased alpha diversity^67^. In particular, only healthy aging mice showed significantly increased Shannon diversity with age. Consistent with previous work^68^, our study also linked aging to an increase in interindividual variation in gut microbial community composition, with interindividual variation being especially high in the normal aging group. This suggested that the unhealthy aging related changes in the gut microbiota are likely stochastic, leading to community instability. Our study also linked FI to several microbial features such as ASVs from *Clostridium sensu stricto, Clostridium XlVa, Enterorhabdus*, and *Phocea massiliensis*. Importantly, we constructed a machine learning model that can predict healthy aging with LOOCV accuracy of 0.773 (17/22) based on these FI related microbial features. And these microbial features may be further driven by CR after 21 months of age. Indeed, we found that some predictive features (e.g., ASVs from *Clostridium sensu stricto* and *Enterorhabdus*) were only identified as differentially abundant taxa at 30 months of age. These findings suggest that key microbial taxa could potentially serve as biomarkers of aging and might contribute to the pathophysiology of aging, although the latter possibility remains to be determined.

We acknowledge the following limitations of this study. First, the sample size of the experimental cohort is relatively small and limited to male mice. Second, 16S rRNA gene sequencing limits our ability to establish associations at the strain level, suggesting that future studies with shotgun metagenomics sequencing will increase resolution. Third, the association between healthy aging and microbial taxa identified in this study does not demonstrate causality. Thus, additional research is needed to validate the mechanism behind these essential findings. Finally, the generalization of the machine learning-based gut microbial signature of aging to other murine cohorts and to humans remains unknown. However, the strengths of the study include a prospective study design, detailed phenotyping of mice, and assessment of accuracy using gut microbial features to predict healthy aging by machine learning model.

In conclusion, we evaluated the impact of age-related changes in gut microbiota on the course of aging in late-life male mice to assess a microbiota signature associated with healthy aging. Our study suggests the possible interaction between specific gut microbiota and aging status, and motivates future work that could establish causality and the potential of future microbiota-targeted interventions to increase healthy aging.

## Methods

### Study population and sample collection

In our study, we only included the mice with basic measurements and biological samples at both 21 and 30 months, resulting 22 C57BL/6 male mice (NIA Aging Colony). Following baseline measurements (body mass, food intake, frailty index and fecal collection), we randomly divided these mice into two diet groups, fed either *ad libitum* (AL, n=14) with standard chow or under mild (15%) calorie restriction (CR, n=8) and followed longitudinally until death. Mice were fed a standard chow based upon AIN-93G (Custom diet #A17101101, Research Diets, New Brunswick, NJ). CR was initiated over a period of two-weeks in a step-down fashion (10% CR, 15% CR) to ensure no loss on mice as they transition to the restricted feeding paradigm. Fecal samples (non-fasted) were collected in the morning (8.30am-11.30am) into sterile tubes and frozen at −80 °C until future analysis.

### The measurement of frailty index

Frailty was measured using the validated 31-parameter mouse clinical frailty index as described previously^36,37^. Briefly, the clinical assessment includes evaluation of the integument, the musculoskeletal system, the vestibulocochlear/auditory systems, ocular and nasal systems, digestive system, urogenital system, respiratory system, signs of discomfort, body mass, and body surface temperature. FI score is continuous from 0-1, with higher values indicating worse frailty^37^.

### Hematology analysis

25μL of whole blood obtained via submandibular bleeding was combined with 1μL of EDTA to prevent clotting. The sample was analyzed using a Hemavet 950 veterinary (Drew Scientific, Miami Lakes, FL) multi-species hematology system using standard settings.

### Estimation of bacterial load by quantitative PCR

To estimate gut bacterial load in our 44 fecal samples, we performed quantitative PCR (qPCR) targeting the 16S rRNA gene using the same primers employed for 16S rRNA gene sequencing (515F and 806R). Briefly, 2□μl of template DNA was combined with 12.5 μl PerfeCTa SYBR Green SuperMix Reaction Mix (QuantaBio, Beverly, MA), 6□μl nuclease-free H2O, and 2.25□μl of each primer. Amplification was performed on a Bio-Rad CFX384 Touch (Bio-Rad, Hercules, CA) in the Bauer Core Facility at Harvard University using the following cycle settings: 95□°C for 10□min, followed by 40 cycles of 95□°C for 15□s, 60□°C for 40□s and 72□°C for 30□s. Reactions were performed in triplicate with the mean value used in statistical analyses. Cycle-threshold values were standardized against a dilution curve of *Escherichia coli* genomic DNA at the following concentrations (ng/μL): 100, 50, 25, 10, 5, 1, 0.5, plus a no-template (negative) control. Bacterial DNA concentrations were normalized to 16S copies/μL, then multiplied by the total extracted DNA volume (50 μL) and divided by the grams of fecal matter utilized in the extraction of template DNA (varied), allowing us to report gut bacterial load as 16S rRNA gene copies per gram of feces.

### DNA isolation and 16S rRNA gene sequencing

Gut microbial DNA was isolated using the DNeasy PowerSoil Pro Kit (Qiagen) and PCR-amplified using barcoded primers targeting the V4 region of the bacterial 16S rRNA gene [515F (GTGYCAGCMGCCGCGGTAA) and 806R (GGACTACNVGGGTWTCTAAT); Integrated DNA Technologies]. The following thermocycler protocol was used: 94°C for 3 min, 35 cycles of 94°C for 45 s, 50°C for 30 s, and 72°C for 90 s, with a final extension at 72°C for 10 min. Triplicate PCR reactions for each sample were pooled and amplification was confirmed by 1.5% gel electrophoresis. 16S rDNA amplicons were cleaned with AmpureXP beads (Agencourt) on a per-sample basis, then quantified using the Quant-iT Picogreen dsDNA Assay Kit (Invitrogen). Amplicons were pooled evenly by DNA content and sequenced on an Illumina HiSeq (1 x 150 bp) at the Bauer Core Facility at Harvard University, generating 234,631 ± 110,737 (mean ± SD) sequences per sample passing filter (range: 75,898 to 391,101) (Table S1).

### Microbiota composition by 16S rRNA gene amplicon analysis

Raw sequencing data was processed and analyzed using Quantitative Insights into Microbial Ecology 2 (QIIME2) pipeline^70^. Single-end sequences were first demultiplexed using the barcode sequences. The sequencing reads were then quality filtered, denoised, and merged using DADA2^71^ to generate the ASV feature table. For taxonomy classification, ASV feature sequences were aligned against SILVA reference database^72^. Additional species level assignment to the NCBI RefSeq^73^ 16S rRNA database supplemented by RDP^74^ was accomplished using the *assignTaxonomy* and *addSpecies* functions of DADA2 R package.

### Statistical analysis

Microbial alpha and beta diversity measures were calculated at the ASV level using the vegan package in R. Principal coordinates analysis (PCoA) plot was generated with Bray-Curtis dissimilarity. The difference in microbiome compositions by different groups were tested by the permutational multivariate analysis of variance (PERMANOVA) using the “adonis” function in the R’s vegan package. All PERMANOVA tests were performed with the 9999 permutations based on the Bray-Curtis dissimilarity. Differences between groups were analyzed using a Wilcoxon–Mann–Whitney test (unpaired) or Wilcoxon signed rank test (paired). The survival probability was computed by the Kaplan-Meier method.

MaAsLin2^44^ (multivariate association with linear model) was used for adjustment of covariates when determining the significance of ASVs contributing to specific hematological variables and FI, while accounting for potentially confounding covariates. The linear mixed models included each mouse’s identifier as random effects and other potential confounders as fixed effects. To be qualified for downstream analyses, a ASV feature needed to be detected at least 10% of samples. The *P* values were then adjusted using the Benjamini–Hochberg FDR method. The microbial features with corrected *q*□value <□0.2 were presented. For differential abundance analysis, we used ANCOM^51^ (analysis of composition of microbiomes), with a Benjamini–Hochberg correction at 5% level of significance, and adjusted for cage, cohort, body mass, and diet. Only the ASVs presented at least 10% of samples were included. To develop a model capable of predicting healthy aging, we implemented Elastic-net (ENET) using R’s caret package. Custom machine learning process was conducted using microbial features at 21 months of age to predict aging status at 30 months of age. We first trained our model with all of microbial features. To further improve the biological plausibility, we then only included the microbial features significantly associated with FI. A total of 14 ASVs were selected based on the *q* value (*q*□<□0.2) from the MaAsLin2 model. Leave-one-out cross-validation (LOOCV) was applied with the *trainControl* function. To further validate our model, a null model was generated with random selected feature (number of features=14, times=100). All statistical analyses were performed using R.

## Data availability

Raw sequencing reads have been deposited in NCBI under accession number PRJNA739980.

## Acknowledgements

We thank Lusheng Huang, Congying Chen, Xu-Wen Wang and Zheng Sun for helpful discussions.

## Author contributions

Y.-Y.L, R.N.C., F.G. and J.R.M. conceived and designed the project. S.J.M., M.R.M. and A.E.K. performed the mice experiments. E.M.V., K.S.C., and R.N.C. performed the 16S rRNA gene sequencing and universal 16S quantitative PCR. S.K. performed all the data analysis. S.K. and Y.-Y.L. wrote the manuscript. All authors analyzed the results and edited the manuscript.

## Supplementary information

**Fig. S1.**
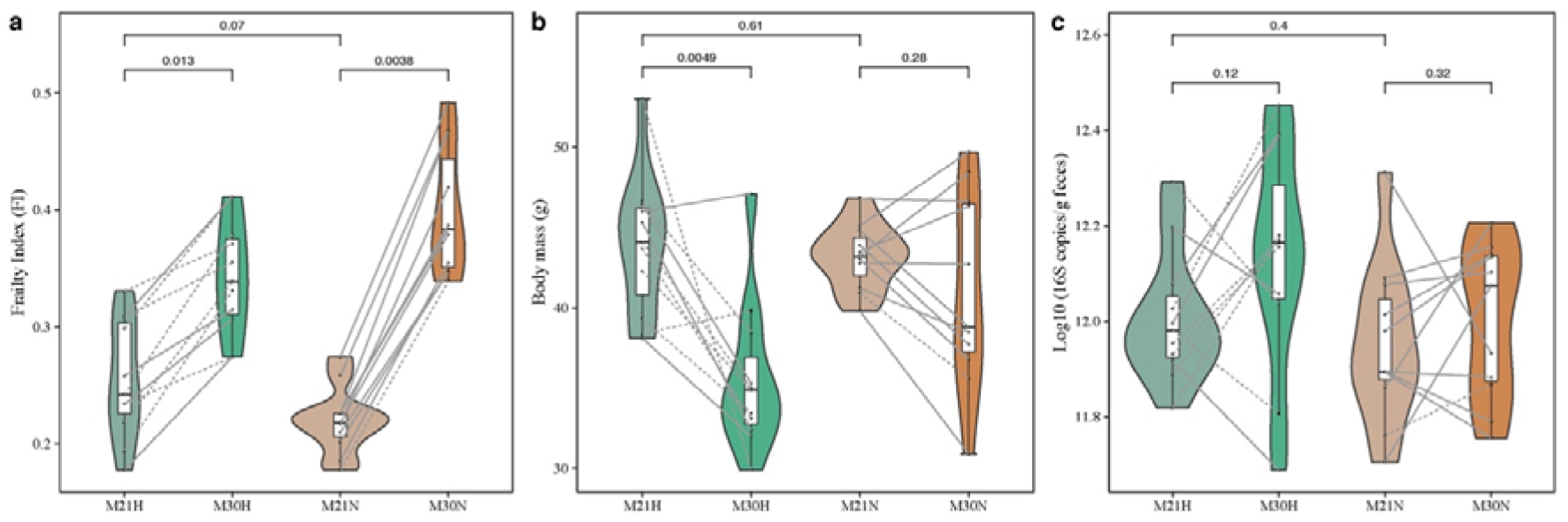
The effects of healthy aging on FI, body mass and total bacterial load. **a**, Frailty index changes with age. Mice were grouped into healthy and normal aging based on the median ΔFI at 30 months of age. **b**, Body mass changes with age. **c**, Total bacterial load changes with age. Points obtained for the same subject from 21 and 30 months of age are joined by solid (AL diet) and dotted (CR diet) lines. *P* value shown the results of Wilcoxon–Mann–Whitney test (unpaired) and Wilcoxon signed rank test (paired).

**Fig. S2.**
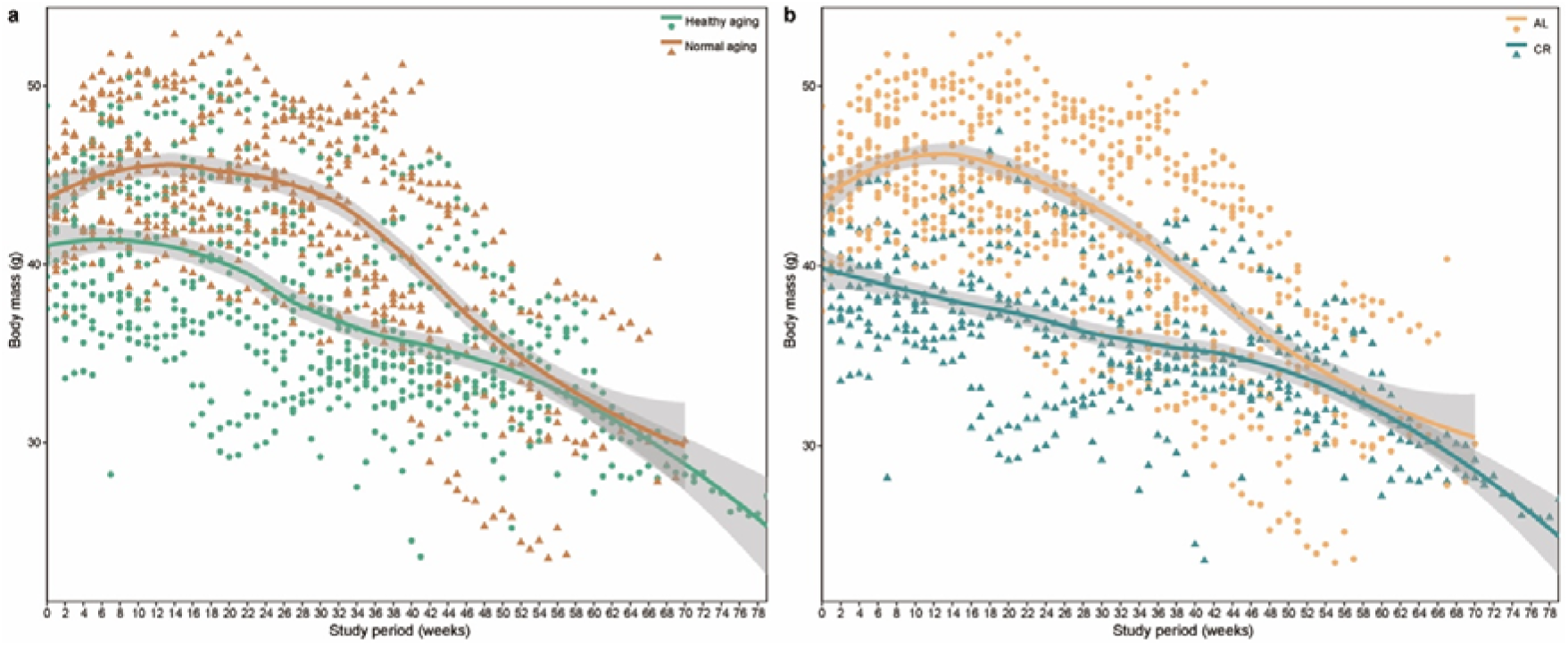
The changes of body mass over time. **a**, Healthy aging versus Normal aging mice. **b**, AL diet versus CR diet. Curves show LOESS fit for the data per category, and shaded areas show 95% confidence intervals for the fit.

**Fig. S3.**
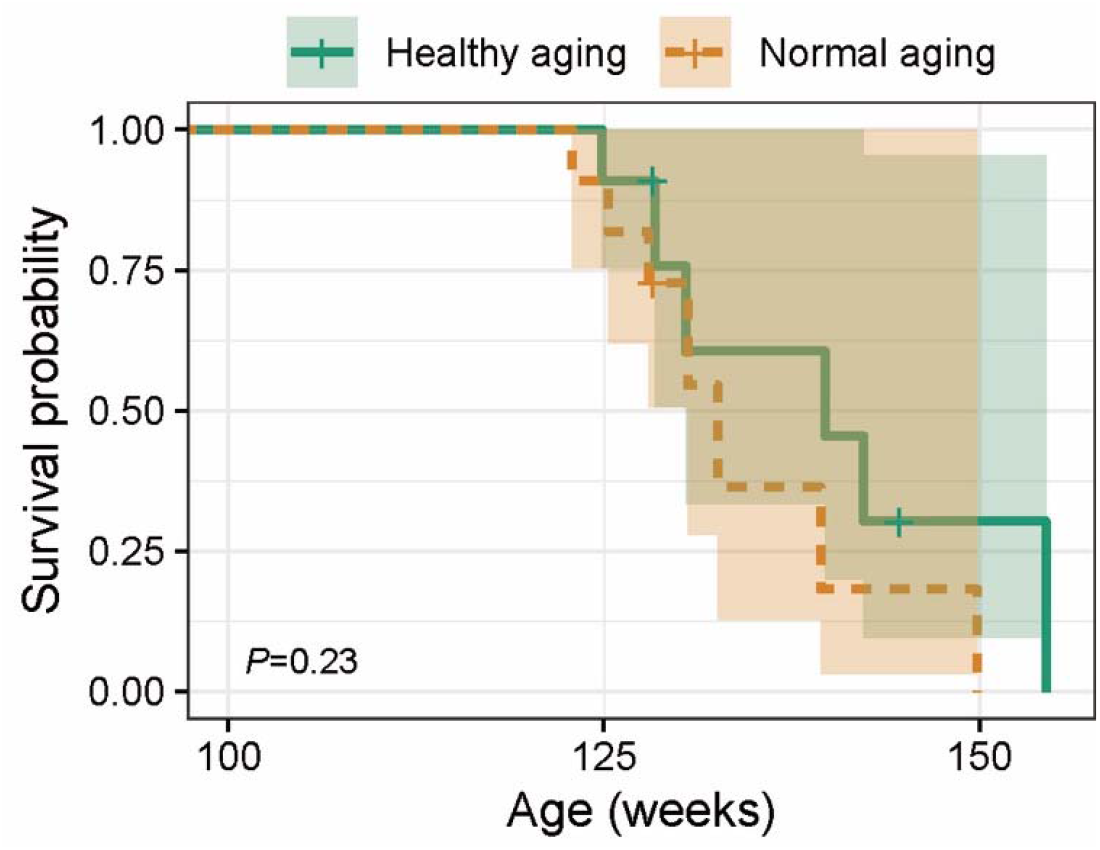
The survival probability was computed by the Kaplan-Meier method. *P* value is the result of log-rank test.

**Fig. S4.**
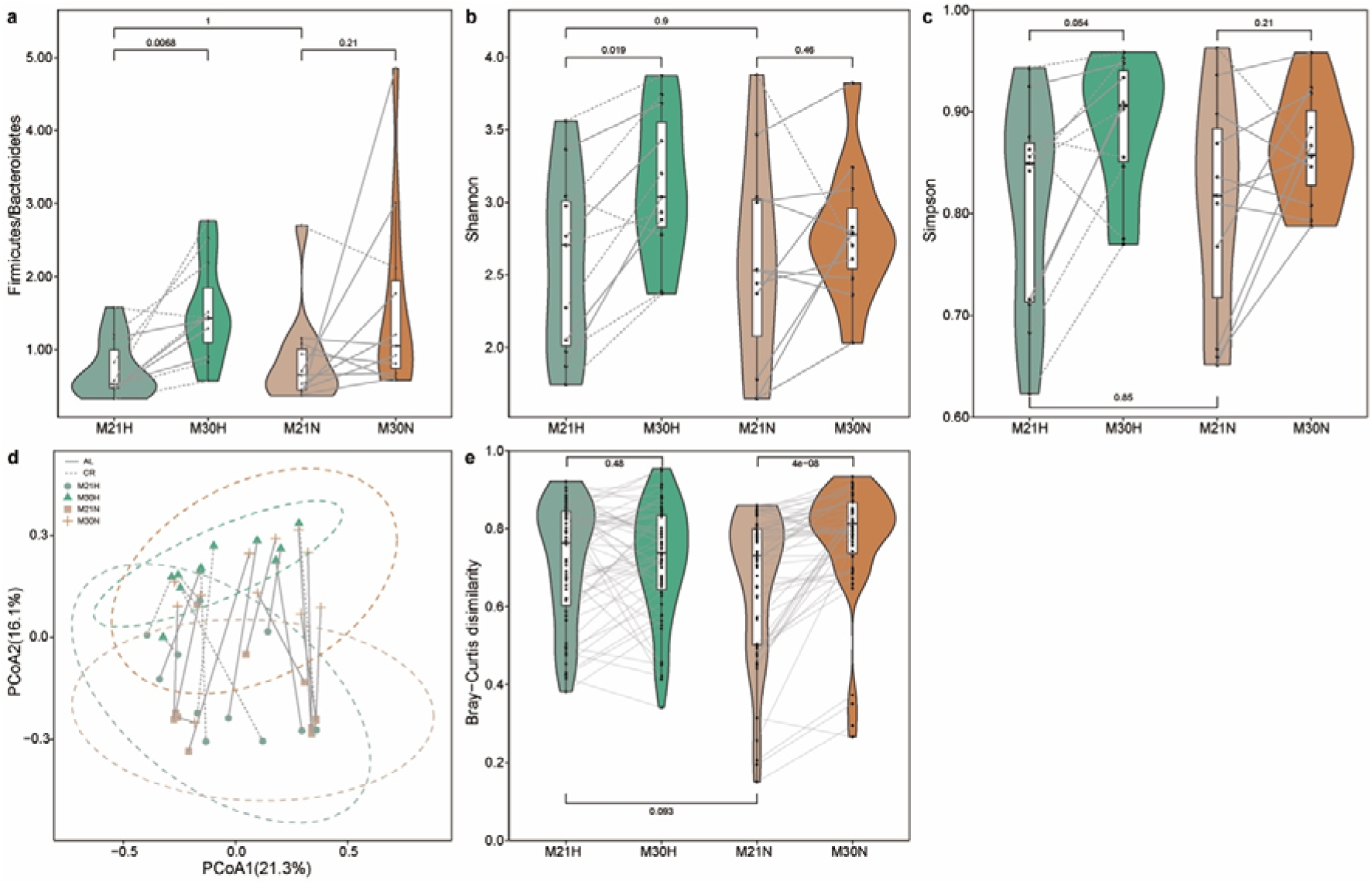
Impact of healthy aging on gut microbial communities. **a**, The ratio of Firmicutes to Bacteroidetes. Alpha diversity using Shannon (**b**) and Simpson (**c**) index. **d**, Beta diversity using Principal Coordinate Analysis (PCoA) of Bray–Curtis dissimilarity. The dotted ellipse borders with color represent the 95% confidence interval. **e**, Boxplot of gut microbiome Bray–Curtis dissimilarity between subjects within each group. Mice were grouped into healthy and normal aging based on the median ΔFI at 30 months of age. Points obtained for the same subject from 21 and 30 months of age in **a**-**d** are joined by solid (AL diet) and dotted (CR diet) lines. Points obtained for the same subject pairs from 21 and 30 months of age in **e** are joined by solid line. *P* value shown are the result of Wilcoxon–Mann–Whitney test (unpaired) and Wilcoxon signed rank test (paired).

**Fig. S5.**
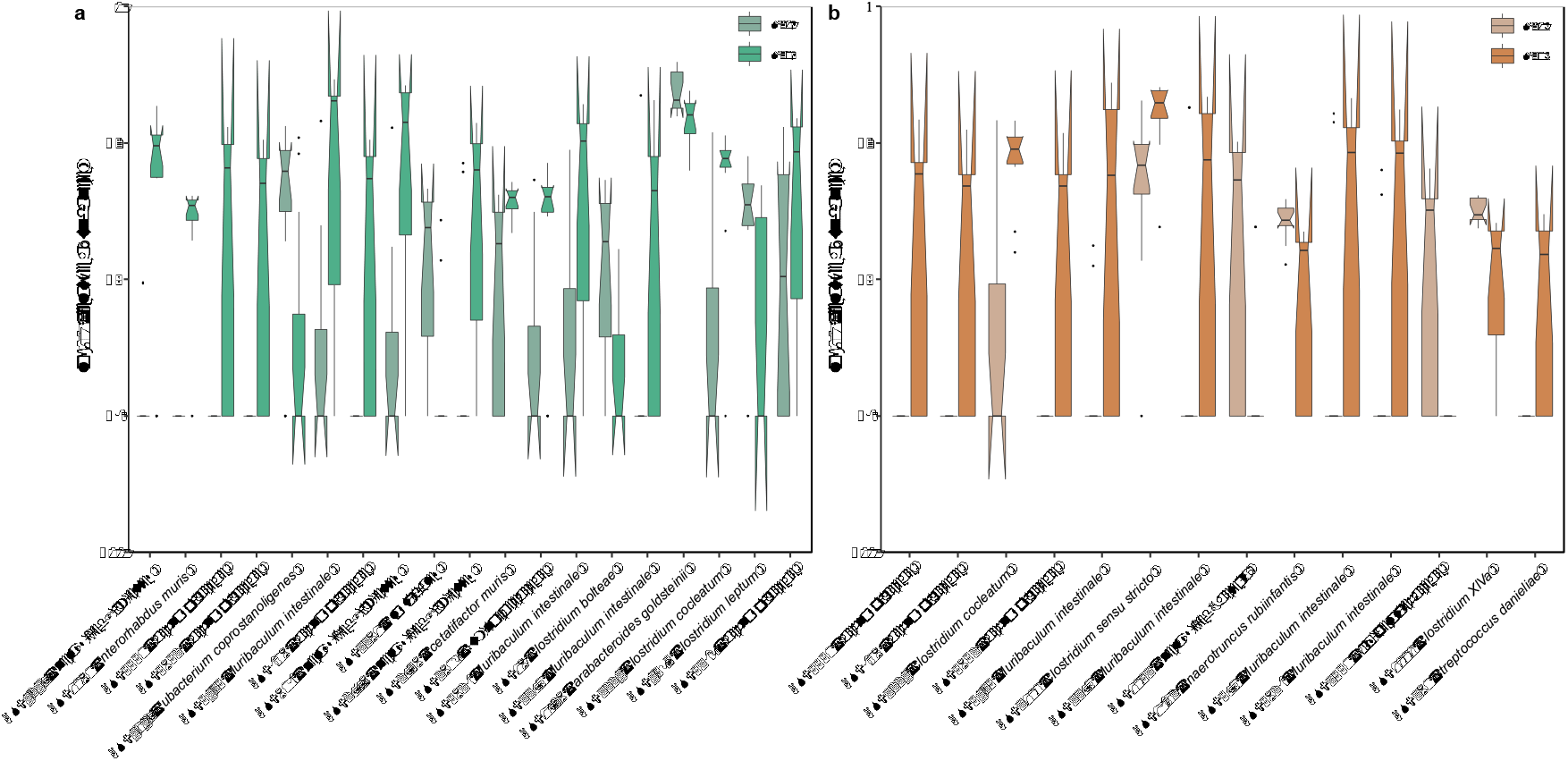
Relative abundance of aging related microbial features in both normal and healthy aging mice. The differential abundant ASVs that differed significantly between 21 and 30months of age for healthy (**a**) and normal (**b**) aging mice identified by analysis of composition of microbiomes (ANCOM). The model was simultaneously adjusted for potential confounders including cage, cohort, diet, and body mass. Mice were grouped into healthy and normal aging based on the median ΔFI at 30 months of age. The top differentially abundant taxa were ranked based on their W statistics (a high “w score” generated by this test indicates the greater likelihood that the null hypothesis can be rejected, indicating the number of times a parameter is significantly different between groups) (from left to right). The relative abundance (%) are plotted on log10 scale. The notches in the boxplots show the 95% confidence interval around the median.

**Fig. S6.**
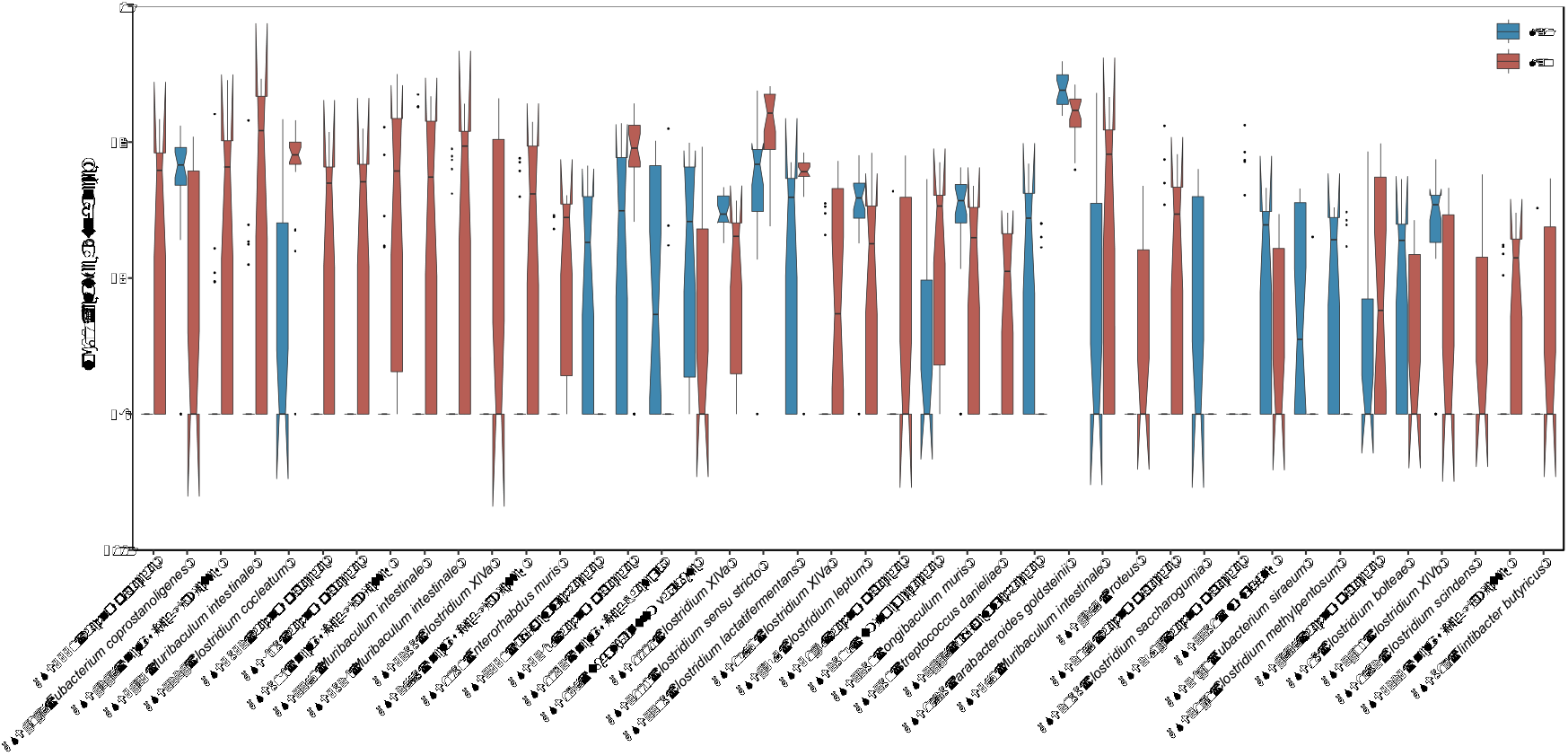
Relative abundance of aging related microbial features. The differential abundant ASVs that differed significantly between 21 and 30months of age identified by ANCOM. The model was simultaneously adjusted for potential confounders including cage, cohort, diet, and body mass. The top differentially abundant taxa were ranked based on their W statistics (a high “w score” generated by this test indicates the greater likelihood that the null hypothesis can be rejected, indicating the number of times a parameter is significantly different between groups) (from left to right). The relative abundance (%) are plotted on log10 scale. The notches in the boxplots show the 95% confidence interval around the median.

**Fig. S7.**
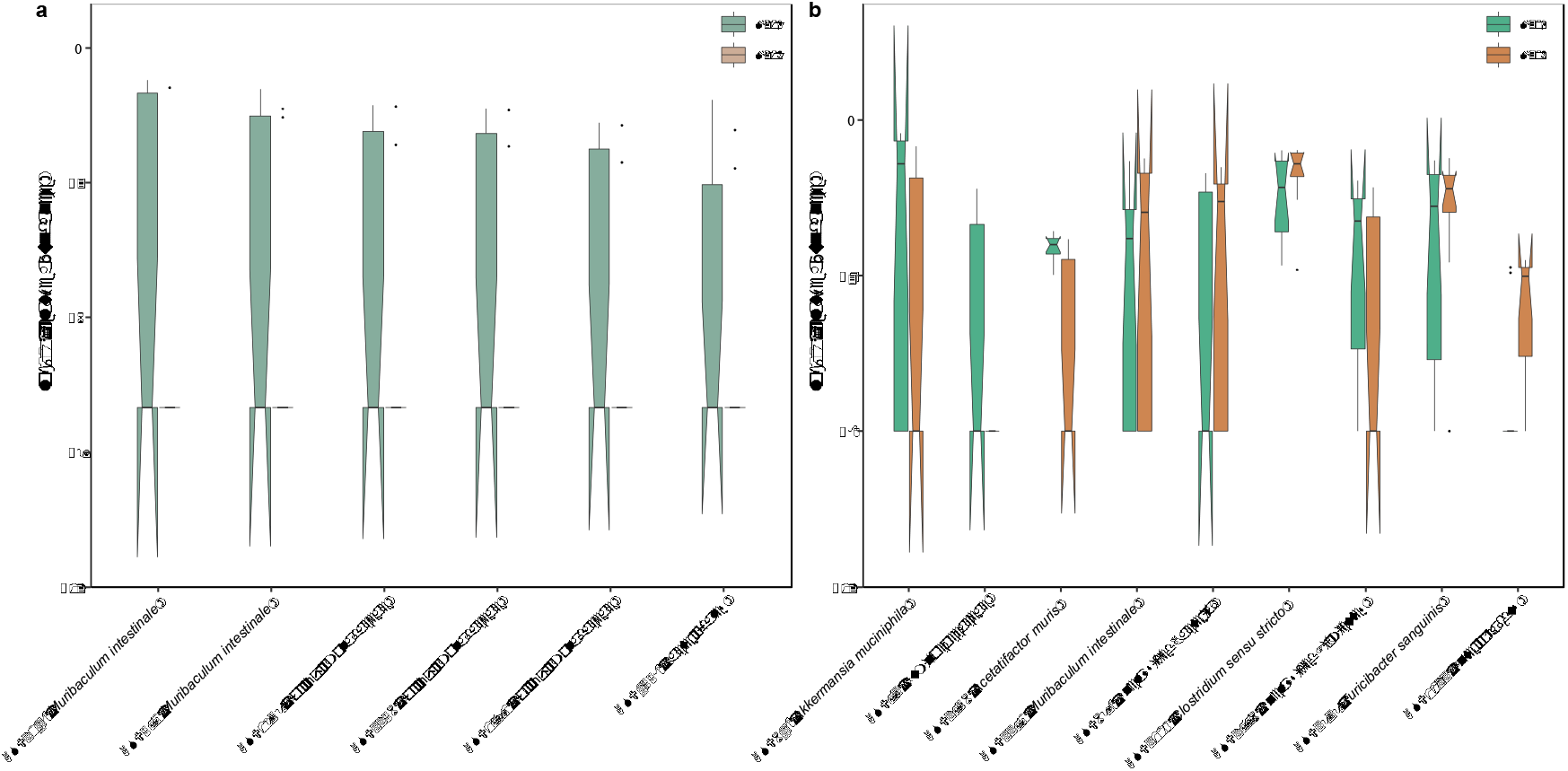
Relative abundance of healthy aging related microbial features. The differential abundant ASVs that differed significantly between healthy and normal aging mice at 21 (**a**) and 30 (**b**) months of ages identified by ANCOM. The model was simultaneously adjusted for potential confounders including cage, cohort, diet, and body mass. Mice were grouped into healthy and normal aging based on the median ΔFI at 30 months of age. The top differentially abundant taxa were ranked based on their W statistics (a high “w score” generated by this test indicates the greater likelihood that the null hypothesis can be rejected, indicating the number of times a parameter is significantly different between groups) (from left to right). The relative abundance (%) are plotted on log10 scale. The notches in the boxplots show the 95% confidence interval around the median.

**Table S1.**
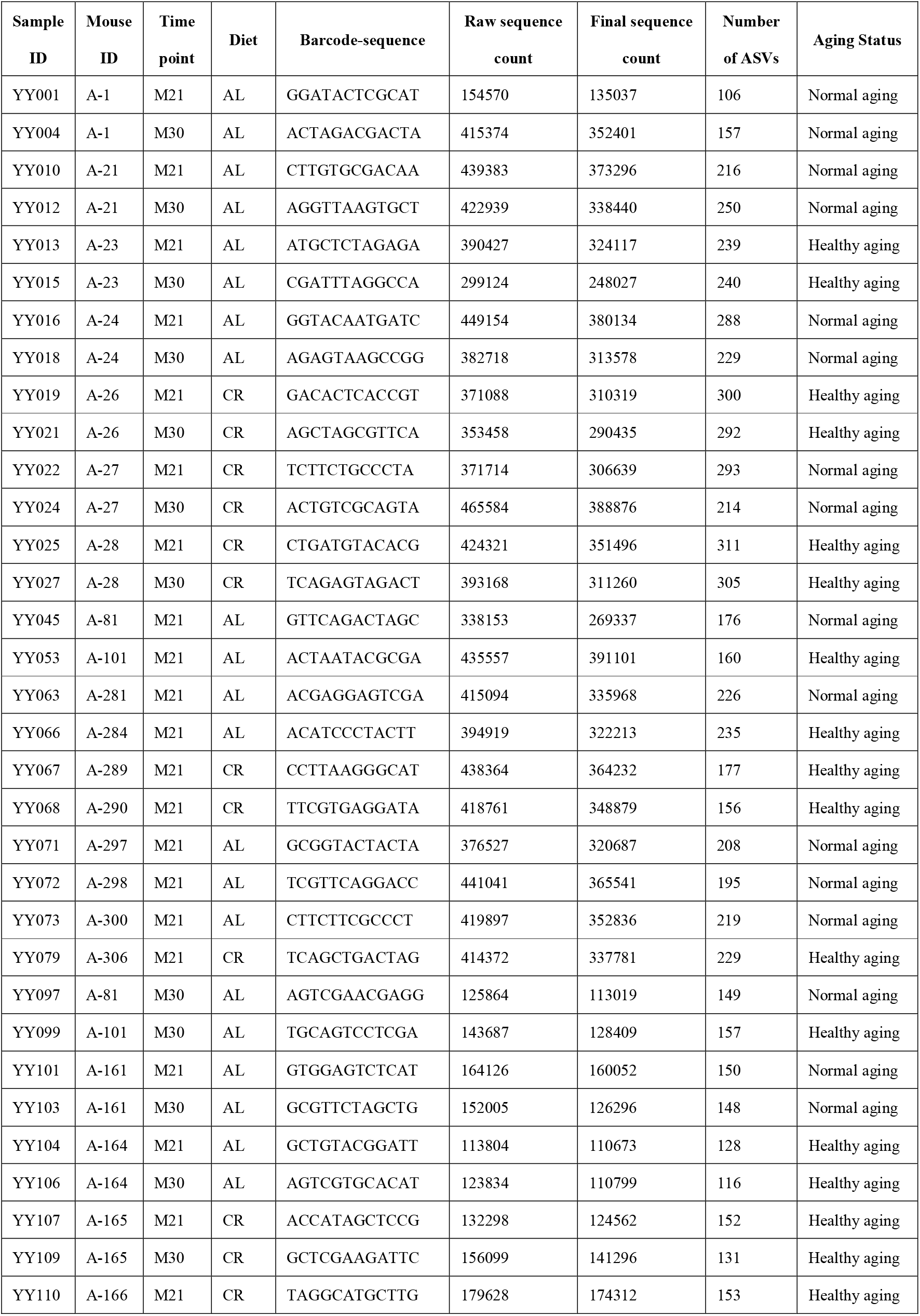
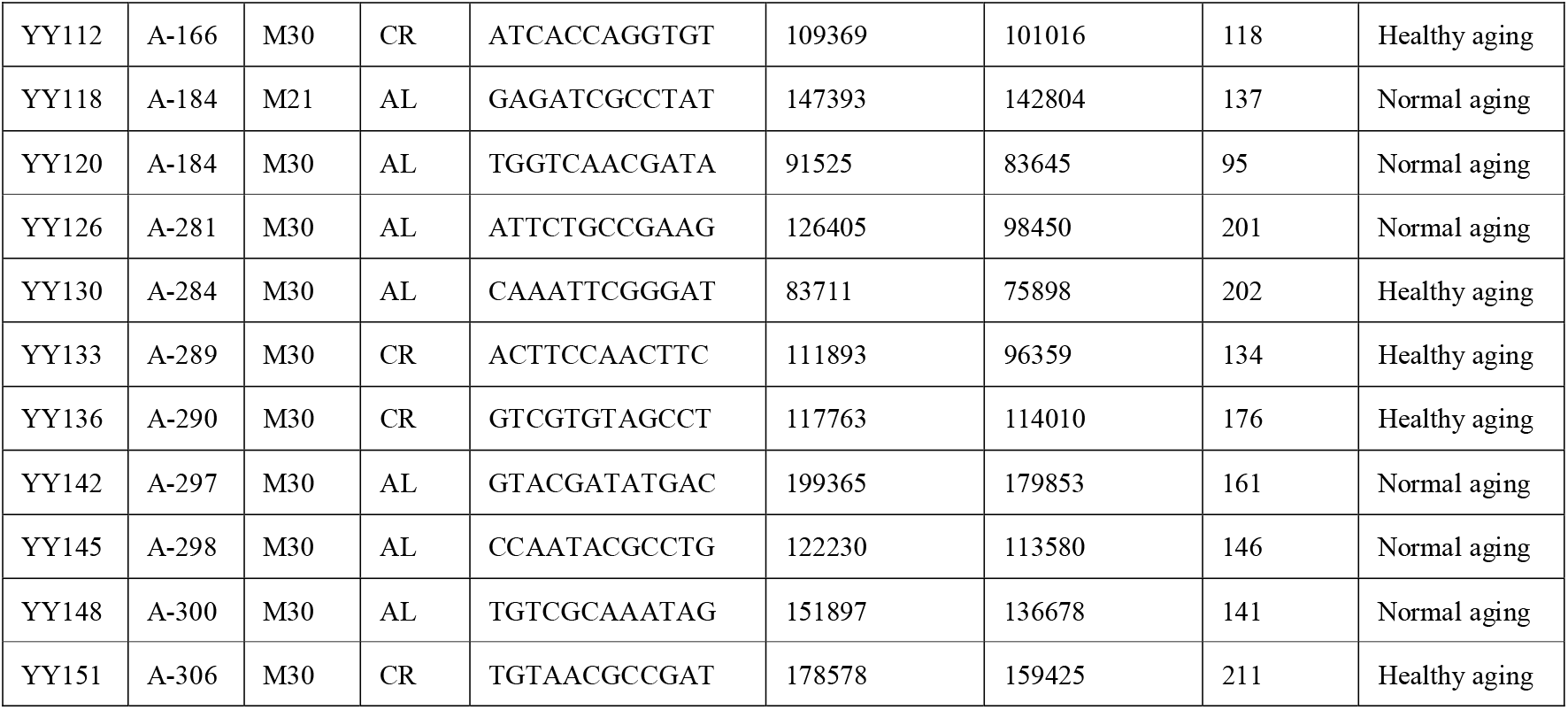
16S rRNA gene sequencing metadata.

**Table S2.**
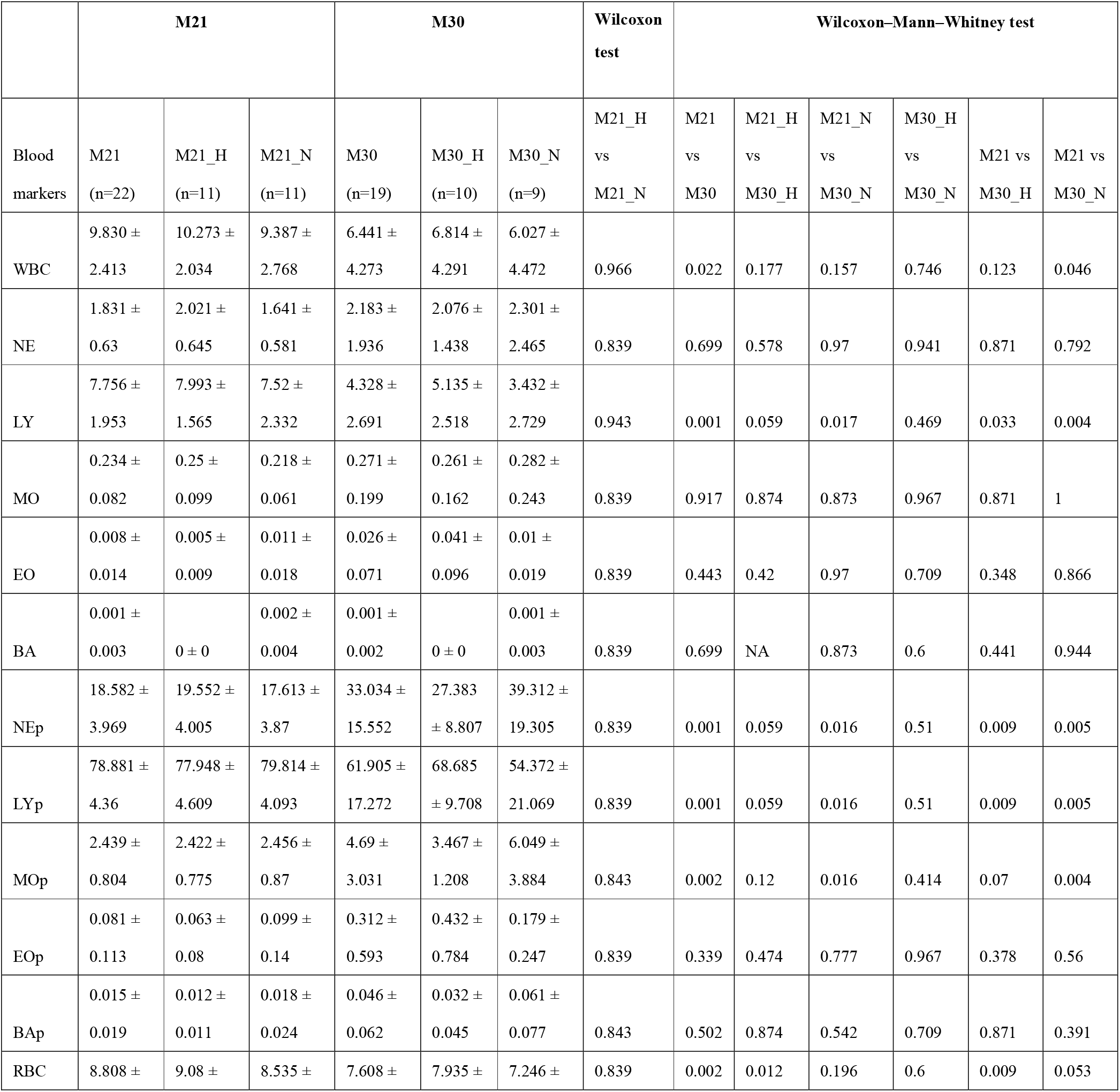
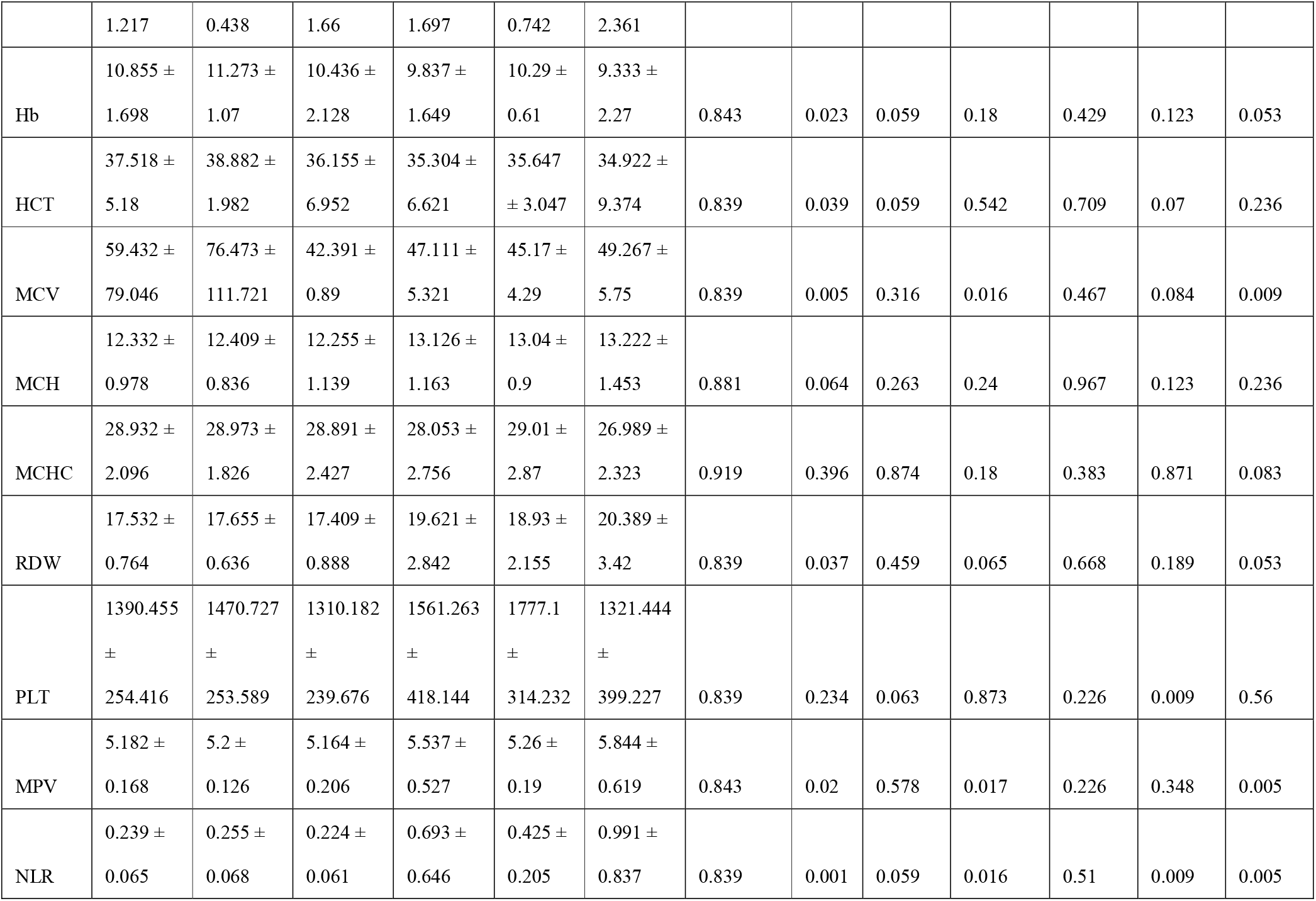
The effect of aging process on blood cells in circulation. The data was shown as mean ± standard deviation. *P* value shown the results of Wilcoxon–Mann–Whitney test (unpaired) and Wilcoxon signed rank test (paired) adjusted using the Benjamini–Hochberg FDR method. WBC: White blood cell, NE: Neutrophils count, LY: Lymphocytes count, MO: Monocytes count, EO: Eosinophils count, BA: Basophils count, NEp: Neutrophils percentage, LYp: Lymphocytes percentage, MOp: Monocytes percentage, EOp: Eosinophils percentage, BA: Basophils percentage, RBC: Red blood cell count, Hb: Hemoglobin, HCT: Hematocrit, MCV: Mean corpuscular volume, MCH: Mean corpuscular hemoglobin, MCHC: Mean corpuscular hemoglobin concentration, RDW: Red cell distribution width, PLT: Platelet count, MPV: Mean platelet volume, NLR: Neutrophils to Lymphocytes ratio.

**Table S3.** The microbial features associated with blood markers identified by MaAsLin2. The relative abundance (%) was shown as mean ± standard deviation.

| ASVs | Taxonomy | Relative abundance (%) |
| --- | --- | --- |
| ASV890 | Ruminococcaceae | 0.004 ± 0.011 |
| ASV806 | Lachnospiraceae | 0.166 ± 0.394 |
| ASV5690 | *Flavonifractor plautii* | 0.242 ± 0.397 |
| ASV5652 | Lachnospiraceae | 0.153 ± 0.355 |
| ASV5625 | Unclassified Firmicutes | 0.054 ± 0.234 |
| ASV5550 | Lachnospiraceae | 0.292 ± 0.671 |
| ASV555 | *Acetatifactor muris* | 0.002 ± 0.005 |
| ASV5396 | *Acetatifactor muris* | 0.025 ± 0.079 |
| ASV5138 | Unclassified Proteobacteria | 0.108 ± 0.321 |
| ASV4558 | Bacteroidales | 0.355 ± 1.284 |
| ASV3949 | *Anaerotruncus* | 0.010 ± 0.018 |
| ASV3897 | Unclassified Bacteria | 0.019 ± 0.056 |
| ASV3729 | *Clostridium aldenense* | 0.003 ± 0.010 |
| ASV3535 | *Muribaculum intestinale* | 0.084 ± 0.226 |
| ASV2973 | *Intestinimonas butyriciproducens* | 0.006 ± 0.016 |
| ASV2878 | Lachnospiraceae | 0.020 ± 0.064 |
| ASV2868 | *Oscillibacter* | 0.007 ± 0.024 |
| ASV2733 | *Clostridium XIVa* | 0.048 ± 0.154 |
| ASV2710 | Unclassified Firmicutes | 0.001 ± 0.002 |
| ASV2261 | Ruminococcaceae | 0.001 ± 0.003 |
| ASV1983 | Ruminococcaceae | 0.043 ± 0.085 |
| ASV1970 | *Clostridium XIVa* | 0.036 ± 0.077 |
| ASV1513 | Lachnospiraceae | 0.002 ± 0.002 |
| ASV1466 | Unclassified Firmicutes | 0.005 ± 0.018 |

**Table S4.** The microbial features associated with Frailty index identified by MaAsLin2. The relative abundance (%) was shown as mean ± standard deviation.

| ASVs | Taxonomy | Relative abundance (%) |
| --- | --- | --- |
| ASV3100 | *Clostridium sensu stricto* | 4.148 ± 5.608 |
| ASV2882 | *Clostridium XIVa* | 0.108 ± 0.133 |
| ASV847 | *Phoceamassiliensis* | 0.004 ± 0.008 |
| ASV338 | Lachnospiraceae | 0.011 ± 0.039 |
| ASV1726 | *Parabacteroides goldsteinii* | 13.017 ± 13.852 |
| ASV5389 | Lachnospiraceae | 1.061 ± 1.702 |
| ASV1123 | *Enterorhabdus* | 0.018 ± 0.025 |
| ASV1101 | *Clostridium XIVa* | 0.025 ± 0.025 |
| ASV807 | Bacteria | 0.001 ± 0.003 |
| ASV742 | Lachnospiraceae | 0.010 ± 0.033 |
| ASV157 | *Subdoligranulum variabile* | 0.131 ± 0.243 |
| ASV232 | Ruminococcaceae | 0.009 ± 0.014 |
| ASV2980 | Lachnospiraceae | 0.101 ± 0.220 |
| ASV466 | Lachnospiraceae | 0.028 ± 0.080 |

**Table S5.** Differentially abundant taxa between 21 and 30 months of age in healthy aging mice detected by ANCOM, adjusted for cage, cohort and diet. For each ASV, the first column represents its taxonomy information, the second column represents its W score and subsequent four columns represent logical indicators of whether it is differentially abundant under a series of cutoffs (0.9, 0.8, 0.7, and 0.6, a prevalence cutoff on the entire set of ASVs). The last two columns denote its relative abundance (%) in each group shown as mean ± standard deviation.

| ASVs | Taxonomy | W_score | detected_0.9 | detected_0.8 | detected_0.7 | detected_0.6 | M21H | M30H |
| --- | --- | --- | --- | --- | --- | --- | --- | --- |
| ASV4247 | Unclassified Firmicutes | 368 | TRUE | TRUE | TRUE | TRUE | 0 ± 0 | 1.357 ± 1.883 |
| ASV1060 | *Enterorhabdus muris* | 352 | TRUE | TRUE | TRUE | TRUE | 0 ± 0 | 0.039 ± 0.023 |
| ASV5550 | Lachnospiraceae | 350 | TRUE | TRUE | TRUE | TRUE | 0 ± 0 | 0.631 ± 0.797 |
| ASV5652 | Lachnospiraceae | 345 | TRUE | TRUE | TRUE | TRUE | 0 ± 0 | 0.331 ± 0.44 |
| ASV4147 | *Eubacterium coprostanoligenes* | 331 | FALSE | TRUE | TRUE | TRUE | 0.622 ± 0.836 | 0.174 ± 0.414 |
| ASV5435 | *Muribaculum intestinale* | 330 | FALSE | TRUE | TRUE | TRUE | 0.275 ± 0.906 | 7.51 ± 8.01 |
| ASV806 | Lachnospiraceae | 326 | FALSE | TRUE | TRUE | TRUE | 0 ± 0 | 0.345 ± 0.446 |
| ASV608 | Unclassified Firmicutes | 324 | FALSE | TRUE | TRUE | TRUE | 0.196 ± 0.648 | 5.741 ± 7.033 |
| ASV3361 | Clostridiales | 277 | FALSE | FALSE | TRUE | TRUE | 0.027 ± 0.032 | 0.002 ± 0.006 |
| ASV2776 | Unclassified Firmicutes | 275 | FALSE | FALSE | TRUE | TRUE | 0.054 ± 0.123 | 0.74 ± 1.023 |
| ASV2756 | *Acetatifactor muris* | 261 | FALSE | FALSE | FALSE | TRUE | 0.017 ± 0.024 | 0.069 ± 0.042 |
| ASV2609 | Ruminococcaceae | 256 | FALSE | FALSE | FALSE | TRUE | 0.018 ± 0.046 | 0.085 ± 0.1 |
| ASV5628 | *Muribaculum intestinale* | 254 | FALSE | FALSE | FALSE | TRUE | 0.145 ± 0.258 | 1.845 ± 2.288 |
| ASV16 | *Clostridium bolteae* | 253 | FALSE | FALSE | FALSE | TRUE | 0.034 ± 0.049 | 0.001 ± 0.002 |
| ASV3370 | *Muribaculum intestinale* | 250 | FALSE | FALSE | FALSE | TRUE | 1.013 ± 3.359 | 1.042 ± 2.593 |
| ASV1726 | *Parabacteroides goldsteinii* | 248 | FALSE | FALSE | FALSE | TRUE | 20.989 ± 20.721 | 5.247 ± 4.615 |
| ASV3224 | *Clostridium cocleatum* | 246 | FALSE | FALSE | FALSE | TRUE | 0.275 ± 0.555 | 0.558 ± 0.461 |
| ASV4595 | *Clostridium leptum* | 234 | FALSE | FALSE | FALSE | TRUE | 0.121 ± 0.169 | 0.024 ± 0.04 |
| ASV5389 | Lachnospiraceae | 229 | FALSE | FALSE | FALSE | TRUE | 0.369 ± 0.731 | 1.192 ± 1.241 |

**Table S6.** Differentially abundant taxa between 21 and 30 months of age in normal aging mice detected by ANCOM, adjusted for cage, cohort and diet. For each ASV, the first column represents its taxonomy information, the second column represents its W score and subsequent four columns represent logical indicators of whether it is differentially abundant under a series of cutoffs (0.9, 0.8, 0.7, and 0.6, a prevalence cutoff on the entire set of ASVs). The last two columns denote its relative abundance (%) in each group shown as mean ± standard deviation.

| ASVs | Taxonomy | W_score | detected_0.9 | detected_0.8 | detected_0.7 | detected_0.6 | M21N | M30N |
| --- | --- | --- | --- | --- | --- | --- | --- | --- |
| ASV5550 | Lachnospiraceae | 334 | TRUE | TRUE | TRUE | TRUE | 0 ± 0 | 0.477 ± 0.938 |
| ASV806 | Lachnospiraceae | 327 | TRUE | TRUE | TRUE | TRUE | 0 ± 0 | 0.283 ± 0.576 |
| ASV3224 | *Clostridium cocleatum* | 327 | TRUE | TRUE | TRUE | TRUE | 0.48 ± 1.005 | 0.955 ± 0.931 |
| ASV5652 | Lachnospiraceae | 326 | TRUE | TRUE | TRUE | TRUE | 0 ± 0 | 0.248 ± 0.481 |
| ASV5435 | *Muribaculum intestinale* | 316 | FALSE | TRUE | TRUE | TRUE | 0.001 ± 0.002 | 4.381 ± 7.109 |
| ASV3100 | *Clostridium sensu stricto* | 296 | FALSE | TRUE | TRUE | TRUE | 1.222 ± 2.458 | 8.285 ± 6.248 |
| ASV3370 | *Muribaculum intestinale* | 285 | FALSE | FALSE | TRUE | TRUE | 1.091 ± 2.427 | 2.785 ± 3.561 |
| ASV1053 | Unclassified Bacteria | 278 | FALSE | FALSE | TRUE | TRUE | 0.331 ± 0.39 | 0.001 ± 0.004 |
| ASV1812 | *Anaerotruncus rubiinfantis* | 263 | FALSE | FALSE | TRUE | TRUE | 0.025 ± 0.017 | 0.004 ± 0.004 |
| ASV570 | *Muribaculum intestinale* | 258 | FALSE | FALSE | TRUE | TRUE | 0.665 ± 1.52 | 1.999 ± 3.191 |
| ASV5628 | *Muribaculum intestinale* | 254 | FALSE | FALSE | TRUE | TRUE | 0.03 ± 0.077 | 1 ± 1.592 |
| ASV3550 | Erysipelotrichaceae | 250 | FALSE | FALSE | FALSE | TRUE | 0.052 ± 0.079 | 0 ± 0 |
| ASV1101 | *Clostridium XIVa* | 238 | FALSE | FALSE | FALSE | TRUE | 0.038 ± 0.023 | 0.007 ± 0.006 |
| ASV360 | *Streptococcus danieliae* | 220 | FALSE | FALSE | FALSE | TRUE | 0 ± 0 | 0.008 ± 0.01 |

**Table S7.** Differentially abundant taxa between 21 and 30 months of age detected by ANCOM, adjusted for cage, cohort and diet. For each ASV, the first column represents its taxonomy information, the second column represents its W score and subsequent four columns represent logical indicators of whether it is differentially abundant under a series of cutoffs (0.9, 0.8, 0.7, and 0.6, a prevalence cutoff on the entire set of ASVs). The last two columns denote its relative abundance (%) in each group shown as mean ± standard deviation.

| ASVs | Taxonomy | W_score | detected_0.9 | detected_0.8 | detected_0.7 | detected_0.6 | M21 | M30 |
| --- | --- | --- | --- | --- | --- | --- | --- | --- |
| ASV5550 | Lachnospiraceae | 375 | TRUE | TRUE | TRUE | TRUE | 0 ± 0 | 0.554 ± 0.853 |
| ASV4147 | *Eubacterium coprostanoligenes* | 372 | TRUE | TRUE | TRUE | TRUE | 0.55 ± 0.661 | 0.178 ± 0.332 |
| ASV4247 | Unclassified Firmicutes | 369 | TRUE | TRUE | TRUE | TRUE | 0.189 ± 0.886 | 1.879 ± 5.061 |
| ASV5435 | *Muribaculum intestinale* | 369 | TRUE | TRUE | TRUE | TRUE | 0.138 ± 0.641 | 5.946 ± 7.562 |
| ASV3224 | *Clostridium cocleatum* | 366 | TRUE | TRUE | TRUE | TRUE | 0.377 ± 0.798 | 0.756 ± 0.745 |
| ASV5652 | Lachnospiraceae | 366 | TRUE | TRUE | TRUE | TRUE | 0 ± 0 | 0.29 ± 0.452 |
| ASV806 | Lachnospiraceae | 365 | TRUE | TRUE | TRUE | TRUE | 0 ± 0 | 0.314 ± 0.504 |
| ASV608 | Unclassified Firmicutes | 360 | TRUE | TRUE | TRUE | TRUE | 0.123 ± 0.466 | 4.633 ± 8.181 |
| ASV3370 | *Muribaculum intestinale* | 357 | TRUE | TRUE | TRUE | TRUE | 1.052 ± 2.86 | 1.914 ± 3.168 |
| ASV5628 | *Muribaculum intestinale* | 354 | TRUE | TRUE | TRUE | TRUE | 0.087 ± 0.195 | 1.422 ± 1.971 |
| ASV5266 | *Clostridium XIVa* | 342 | FALSE | TRUE | TRUE | TRUE | 0 ± 0 | 1.084 ± 2.245 |
| ASV2776 | Unclassified Firmicutes | 341 | FALSE | TRUE | TRUE | TRUE | 0.05 ± 0.125 | 0.534 ± 0.842 |
| ASV1060 | *Enterorhabdus muris* | 340 | FALSE | TRUE | TRUE | TRUE | 0.003 ± 0.007 | 0.025 ± 0.023 |
| ASV3550 | Erysipelotrichaceae | 332 | FALSE | TRUE | TRUE | TRUE | 0.056 ± 0.093 | 0 ± 0 |
| ASV5389 | Lachnospiraceae | 327 | FALSE | TRUE | TRUE | TRUE | 0.432 ± 0.738 | 1.69 ± 2.134 |
| ASV1053 | Unclassified Bacteria | 324 | FALSE | TRUE | TRUE | TRUE | 0.224 ± 0.348 | 0.09 ± 0.417 |
| ASV157 | *Subdoligranulum variable* | 318 | FALSE | TRUE | TRUE | TRUE | 0.196 ± 0.284 | 0.066 ± 0.176 |
| ASV1101 | *Clostridium XIVa* | 318 | FALSE | TRUE | TRUE | TRUE | 0.039 ± 0.027 | 0.012 ± 0.013 |
| ASV3100 | *Clostridium sensu stricto* | 318 | FALSE | TRUE | TRUE | TRUE | 1.776 ± 3.757 | 6.518 ± 6.203 |
| ASV3306 | *Clostridium lactatifermentans* | 316 | FALSE | TRUE | TRUE | TRUE | 0.098 ± 0.112 | 0.261 ± 0.154 |
| ASV1970 | *Clostridium XIVa* | 312 | FALSE | TRUE | TRUE | TRUE | 0.005 ± 0.013 | 0.071 ± 0.103 |
| ASV4595 | *Clostridium leptum* | 310 | FALSE | TRUE | TRUE | TRUE | 0.102 ± 0.126 | 0.055 ± 0.129 |
| ASV5149 | Lachnospiraceae | 310 | FALSE | TRUE | TRUE | TRUE | 0.004 ± 0.017 | 0.046 ± 0.109 |
| ASV2609 | Ruminococcaceae | 308 | FALSE | TRUE | TRUE | TRUE | 0.02 ± 0.043 | 0.058 ± 0.078 |
| ASV3260 | *Longibaculum muris* | 308 | FALSE | TRUE | TRUE | TRUE | 0.068 ± 0.07 | 0.025 ± 0.033 |
| ASV360 | *Streptococcus danieliae* | 305 | FALSE | FALSE | TRUE | TRUE | 0 ± 0 | 0.006 ± 0.009 |
| ASV3447 | Erysipelotrichaceae | 300 | FALSE | FALSE | TRUE | TRUE | 0.058 ± 0.096 | 0.002 ± 0.005 |
| ASV1726 | *Parabacteroides* | 299 | FALSE | FALSE | TRUE | TRUE | 19.963 ± | 6.07 ± 4.97 |
|  | *goldsteinii* |  |  |  |  |  | 16.342 |  |
| ASV570 | *Muribaculum intestinale* | 292 | FALSE | FALSE | TRUE | TRUE | $1.52 \pm 3.134$ | $1.592 \pm 2.524$ |
| ASV4275 | *Proteus* | 284 | FALSE | FALSE | TRUE | TRUE | $0 \pm 0$ | $0.01 \pm 0.025$ |
| ASV2075 | Lachnospiraceae | 283 | FALSE | FALSE | TRUE | TRUE | $0.131 \pm 0.492$ | $0.097 \pm 0.154$ |
| ASV2066 | *Clostridium*<br>*saccharogumia* | 280 | FALSE | FALSE | TRUE | TRUE | $0.05 \pm 0.084$ | $0 \pm 0$ |
| ASV2904 | Lachnospiraceae | 278 | FALSE | FALSE | TRUE | TRUE | $0 \pm 0$ | $0.174 \pm 0.52$ |
| ASV3361 | Clostridiales | 274 | FALSE | FALSE | TRUE | TRUE | $0.021 \pm 0.025$ | $0.004 \pm 0.007$ |
| ASV3840 | *Eubacterium siraeum* | 272 | FALSE | FALSE | TRUE | TRUE | $0.023 \pm 0.033$ | $0 \pm 0.002$ |
| ASV3141 | *Clostridium*<br>*methylopentosum* | 270 | FALSE | FALSE | TRUE | TRUE | $0.012 \pm 0.013$ | $0.003 \pm 0.008$ |
| ASV4737 | Lachnospiraceae | 267 | FALSE | FALSE | FALSE | TRUE | $0.059 \pm 0.173$ | $0.159 \pm 0.275$ |
| ASV16 | *Clostridium bolteae* | 253 | FALSE | FALSE | FALSE | TRUE | $0.02 \pm 0.037$ | $0.002 \pm 0.005$ |
| ASV3400 | *Clostridium XIVb* | 253 | FALSE | FALSE | FALSE | TRUE | $0.068 \pm 0.098$ | $0.017 \pm 0.026$ |
| ASV1762 | *Clostridium scindens* | 249 | FALSE | FALSE | FALSE | TRUE | $0 \pm 0$ | $0.022 \pm 0.047$ |
| ASV5225 | Unclassified Firmicutes | 244 | FALSE | FALSE | FALSE | TRUE | $0.001 \pm 0.002$ | $0.006 \pm 0.007$ |
| ASV613 | *Flintibacter butyricus* | 232 | FALSE | FALSE | FALSE | TRUE | $0.002 \pm 0.007$ | $0.015 \pm 0.036$ |

**Table S8.** Differentially abundant taxa between healthy and normal aging mice at 21 months of age detected by ANCOM, adjusted for cage, cohort and diet. For each ASV, the first column represents its taxonomy information, the second column represents its W score and subsequent four columns represent logical indicators of whether it is differentially abundant under a series of cutoffs (0.9, 0.8, 0.7, and 0.6, a prevalence cutoff on the entire set of ASVs). The last two columns denote its relative abundance (%) in each group shown as mean ± standard deviation.

| ASVs | Taxonomy | W_score | detected_<br>0.9 | detected_<br>0.8 | detected_<br>0.7 | detected_<br>0.6 | Healthy aging<br>(M21H) | Normal aging<br>(M21N) |
| --- | --- | --- | --- | --- | --- | --- | --- | --- |
| ASV2048 | *Muribaculum intestinale* | 366 | TRUE | TRUE | TRUE | TRUE | 5.129 ± 7.579 | 1.2 ± 3.98 |
| ASV570 | *Muribaculum intestinale* | 289 | FALSE | FALSE | TRUE | TRUE | 2.375 ± 4.087 | 0.665 ± 1.52 |
| ASV1959 | Porphyromonadaceae | 268 | FALSE | FALSE | TRUE | TRUE | 0.982 ± 1.702 | 0.513 ± 1.483 |
| ASV3256 | Porphyromonadaceae | 260 | FALSE | FALSE | FALSE | TRUE | 0.841 ± 1.441 | 0.44 ± 1.258 |
| ASV1791 | Porphyromonadaceae | 238 | FALSE | FALSE | FALSE | TRUE | 0.395 ± 0.688 | 0.202 ± 0.58 |
| ASV4558 | Bacteroidales | 232 | FALSE | FALSE | FALSE | TRUE | 1.063 ± 2.401 | 0.156 ± 0.452 |

**Table S9.** Differentially abundant taxa between healthy and normal aging mice at 30 months of age detected by ANCOM, adjusted for cage, cohort and diet. For each ASV, the first column represents its taxonomy information, the second column represents its W score and subsequent four columns represent logical indicators of whether it is differentially abundant under a series of cutoffs (0.9, 0.8, 0.7, and 0.6, a prevalence cutoff on the entire set of ASVs). The last two columns denote its relative abundance (%) in each group shown as mean ± standard deviation.

| ASVs | Taxonomy | W_score | detected_0.9 | detected_0.8 | detected_0.7 | detected_0.6 | Healthy aging (M30H) | Normal aging (M30N) |
| --- | --- | --- | --- | --- | --- | --- | --- | --- |
| ASV648 | *Akkermansia muciniphila* | 323 | TRUE | TRUE | TRUE | TRUE | 15.487 ± 18.623 | 3.812 ± 6.979 |
| ASV73 | Ruminococcaceae | 300 | FALSE | TRUE | TRUE | TRUE | 0.298 ± 0.566 | 0 ± 0 |
| ASV2756 | *Acetatifactor muris* | 270 | FALSE | FALSE | TRUE | TRUE | 0.069 ± 0.042 | 0.02 ± 0.033 |
| ASV3370 | *Muribaculum intestinale* | 258 | FALSE | FALSE | TRUE | TRUE | 1.042 ± 2.593 | 2.785 ± 3.561 |
| ASV698 | Unclassified Bacteria | 253 | FALSE | FALSE | TRUE | TRUE | 0.935 ± 1.527 | 1.547 ± 2.031 |
| ASV3100 | Clostridium sensu stricto | 248 | FALSE | FALSE | TRUE | TRUE | 4.75 ± 5.907 | 8.285 ± 6.248 |
| ASV2776 | Unclassified Firmicutes | 228 | FALSE | FALSE | FALSE | TRUE | 0.74 ± 1.023 | 0.329 ± 0.591 |
| ASV3939 | *Turicibacter sanguinis* | 218 | FALSE | FALSE | FALSE | TRUE | 2.442 ± 3.116 | 2.59 ± 3.045 |
| ASV1123 | *Enterorhabdus* | 216 | FALSE | FALSE | FALSE | TRUE | 0.003 ± 0.006 | 0.011 ± 0.009 |

